# Single-spore germination analyses reveal that calcium released during *Clostridioides difficile* germination functions in a feed-forward loop

**DOI:** 10.1101/2022.11.11.516080

**Authors:** John W. Ribis, Luana Melo, Shailab Shrestha, David Giacalone, Enrique E. Rodriguez, Aimee Shen, Amy Rohlfing

## Abstract

*Clostridioides difficile* infections begin when its metabolically dormant spores germinate in response to sensing bile acid germinants alongside amino acid and divalent cation co-germinants in the small intestine. While bile acid germinants are essential for *C. difficile* spore germination, it is currently unclear whether both co-germinant signals are required. One model proposes that divalent cations, particularly Ca^2+^, are essential for inducing germination, while another proposes that either co-germinant class can induce germination. The former model is based on the finding that spores defective in releasing large stores of internal Ca^2+^ in the form of calcium dipicolinic acid (CaDPA) cannot germinate when germination is induced with bile acid germinant and amino acid co-germinant alone. However, since the reduced optical density of CaDPA-less spores makes it difficult to accurately measure their germination, we developed a novel automated, time-lapse microscopy-based germination assay to analyze CaDPA mutant germination at the single-spore level. Using this assay, we found that CaDPA mutant spores germinate in the presence of amino acid co-germinant and bile acid germinant. Higher levels of amino acid co-germinants are nevertheless required to induce CaDPA mutant spores to germinate relative to WT spores because CaDPA released by WT spores during germination can function in a feedforward loop to potentiate the germination of other spores within the population. Collectively, these data indicate that Ca^2+^ is not essential for inducing *C. difficile* spore germination because amino acid and Ca^2+^ co-germinant signals are sensed by parallel signaling pathways.

**Importance:** *Clostridioides difficile* spore germination is essential for this major nosocomial pathogen to initiate infection. *C. difficile* spores germinate in response to sensing bile acid germinant signals alongside co-germinant signals. There are two classes of co-germinant signals: Ca^2+^ and amino acids. Prior work suggested that Ca^2+^ is essential for *C. difficile* spore germination based on bulk population analyses of germinating CaDPA mutant spores. Since these assays rely on optical density to measure spore germination and the optical density of CaDPA mutant spores is reduced relative to WT spores, this bulk assay is limited in its capacity to analyze germination. To overcome this limitation, we developed an automated image analysis pipeline to monitor *C. difficile* spore germination using time-lapse microscopy. With this analysis pipeline, we demonstrate that, although Ca^2+^ is dispensable for inducing *C. difficile* spore germination, CaDPA can function in a feedforward loop to potentiate the germination of neighboring spores.

## Introduction

*Clostridioides difficile* is the leading cause of nosocomial infections and antibiotic-associated gastroenteritis worldwide. *C. difficile* causes ∼450,000 infections and $5 billion dollars in associated medical costs annually in the United States (1, 2). Prior antibiotic exposure can render otherwise healthy individuals susceptible to *C. difficile* infections (CDI) because a diverse gut flora provides colonization resistance against CDI (3, 4). Once individuals become infected with *C. difficile*, they are highly susceptible to subsequent infections, with about one in five individuals experiencing at least one relapse or recurrence of CDI (2). This high incidence of repeat infections is a major contributor to CDI morbidity and mortality.

Both the vegetative and metabolically dormant spore forms of *C. difficile* contribute to the high rates of *C. difficile* disease recurrence. Vegetative *C. difficile* cells produce the glucosylating toxins, TcdA and TcdB, that induce inflammation and perpetuate gut dysbiosis (5), while its aerotolerant spores allow this obligate anaerobe to transmit disease (6). *C. difficile* spores can survive the antibiotic regimens used to treat *C. difficile* infections because they are metabolically dormant, and their resistance properties also allow them to survive exposure to commonly used disinfectants. As a result, spores serve as an environment reservoir for seeding future infections (7-10).

*C. difficile* infections begin when spores ingested by a susceptible individual germinate, i.e. exit dormancy, in response to specific signals sensed in the small intestine (11-13). Since germination is essential for *C. difficile* to initiate infection, its germination proteins represent promising targets for therapies designed to prevent CDI (13, 14). These proteins are likely to be selective targets because *C. difficile* uses a unique signaling pathway to initiate spore germination relative to previously characterized spore-forming bacteria such as *Bacillus subtilis* and *Clostridium perfringens* (15, 16). Almost all spore-formers use transmembrane Ger-type germinant receptors found in the inner spore membrane (17) to sense nutrient signals (e.g. amino acids and sugars) and induce germination (16, 18). In contrast, *C. difficile* is one of only two spore-formers known to lack Ger-type germinant receptors (19). Instead, *C. difficile* uses the soluble pseudoprotease CspC to sense cholate-derived bile acid germinants (20), which are found exclusively in the gastrointestinal tract of metazoans (21, 22).

Notably, bile acid germinants alone cannot induce *C. difficile* spore germination because they must be combined with co-germinant signal(s) (11). These co-germinant signals belong to two classes: amino acids and divalent cations (23, 24). Glycine is the most potent amino acid co-germinant (23, 25), and Ca^2+^ is the most potent divalent cation co-germinant (24, 26). While the chemical nature of these co-germinant signals has been well characterized, the molecular mechanism by which they are physically sensed in *C. difficile* remains unclear. This is because mutations in two separate genes alter co-germinant sensing. Certain mutations in the *cspBA* gene, which encodes a fusion of the CspB protease to the CspA pseudoprotease (27), allow *C. difficile* spores to germinate independent of co-germinant signals, namely in the presence of bile acids alone (28). Since these mutations alter the site of interdomain processing in CspBA, which changes the N-terminus of CspA (29), the CspA pseudoprotease has been proposed to function as the receptor for both amino acid and Ca^2+^ co-germinants (28). While specific CspA mutations allow *C. difficile* to bypass the need for co-germinants (28), specific amino acid substitutions in CspC, the bile acid germinant receptor, can also alter the sensitivity of *C. difficile* spores to distinct classes of co-germinants (30). These observations indicate that CspC plays a key role in integrating signals from bile acid germinants *and* the two distinct co-germinant classes via an unknown mechanism.

In response to sensing germinants and co-germinants, CspC and CspA somehow activate the CspB protease (11, 13). This allows activated CspB to proteolytically activate the cortex lytic enzyme, SleC (27), which then degrades the cortex, a protective layer of modified peptidoglycan critical for maintaining the metabolic dormancy of spores. Degradation of the cortex activates the mechanosensitive SpoVAC channel in the inner spore membrane (31-33), leading to the release of large internal stores of the spore-specific small molecule, calcium dipicolinic acid (CaDPA), from the spore core (34). This allows water to enter the partially dehydrated spore core, leading to the resumption of metabolism and subsequent outgrowth of the germinating spore into a vegetative cell.

Even though several proteins involved in transducing germinant and co-germinant signals have been identified, there is still considerable debate regarding how *C. difficile* senses the two distinct classes of co-germinant signals. Kochan *et al*. first identified that Ca^2+^ is a potent co-germinant for *C. difficile* spores (24). Kochan *et al.* also showed that a CaDPA-less mutant cannot germinate in the presence of bile acid germinant and amino acid co-germinant alone and that the Ca^2+^ chelator, EGTA, can inhibit germination induced under these conditions (24).

These findings led them to propose that Ca^2+^ is absolutely required for *C. difficile* spore germination, perhaps because Ca^2+^ is used as a co-factor by a key germination protein like CspB (12, 24). According to this ‘lock-and-key’ model, the two classes of co-germinants function sequentially in the same signaling pathway, but only Ca^2+^ is absolutely required as a co-germinant. The alternative model is that either co-germinant class can potentiate bile acid-induced germination (11, 13) such that there are parallel mechanisms for sensing the chemically distinct amino acid and Ca^2+^ co-germinant signals. This model is supported by the identification of specific CspC amino acid substitutions that sensitize spores to either amino acid or divalent cation co-germinants (30).

In this study, we distinguish between these models by rigorously assessing the requirement for Ca^2+^ during *C. difficile* spore germination in a panel of CaDPA-less mutant spores. As CaDPA-less spores are less optically dense than spores containing CaDPA (35), it can be challenging to use traditional optical density-based methods to measure the germination of bulk populations. To overcome these limitations, we developed a novel time-lapse microscopy-based method to simultaneously measure the germination kinetics of hundreds of spores in a given experiment using automated image analysis. With this single-spore germination assay, we determined that germination induced by glycine co-germinant and bile acid germinant (i.e. in the absence of added Ca^2+^) is less efficient in CaDPA-less spores relative to WT. We further showed that this difference in germination behavior is because endogenous Ca^2+^ released from WT spores in the form of CaDPA during germination can function in a positive feedback loop to promote the germination of neighboring spores. Taken together, our data strongly suggest that parallel pathways transduce signals from amino acid and Ca^2+^ co-germinant classes and support a model in which CspC and CspA function in concert to integrate signals from multiple environmental sources to initiate infection.

## Results

### Phenotypic characterization of CaDPA mutant spores

The proposal that Ca^2+^ is absolutely required for *C. difficile* spore germination is largely based on the finding that spores lacking CaDPA fail to germinate when germination is induced by bile acid germinant taurocholate (TA) and glycine co-germinant alone (24). Since the mutant spores in this study lack CaDPA due to the loss of a previously uncharacterized putative ATP-binding protein, CD3298, we sought to analyze the germination phenotypes of additional CaDPA mutant spores lacking known regulators of either CaDPA synthesis or transport into the spore core. Namely, we deleted *dpaAB*, *spoVV*, and *spoVAC* using the *pyrE*-based allelic exchange system from the 630Δ*erm*Δ*pyrE* strain background. *dpaAB* encodes the enzyme complex that synthesizes CaDPA (**Fig 1A**, (35, 36)), while SpoVV and SpoVAC are critical in *B. subtilis* for transporting CaDPA across the outer and then inner forespore membranes, respectively (37, 38) (**Fig 1A**). SpoVV function was first established in *B. subtilis* (37), whereas SpoVAC has been shown to transport CaDPA into the spore core in several spore-formers, including *C. difficile* (32, 35, 39-43). CD3298 may act as an accessory protein for the SpoVV or the SpoVAC transporter, although CD3298 lacks a signal sequence ((24), **Fig 1A**).

**Figure 1.**
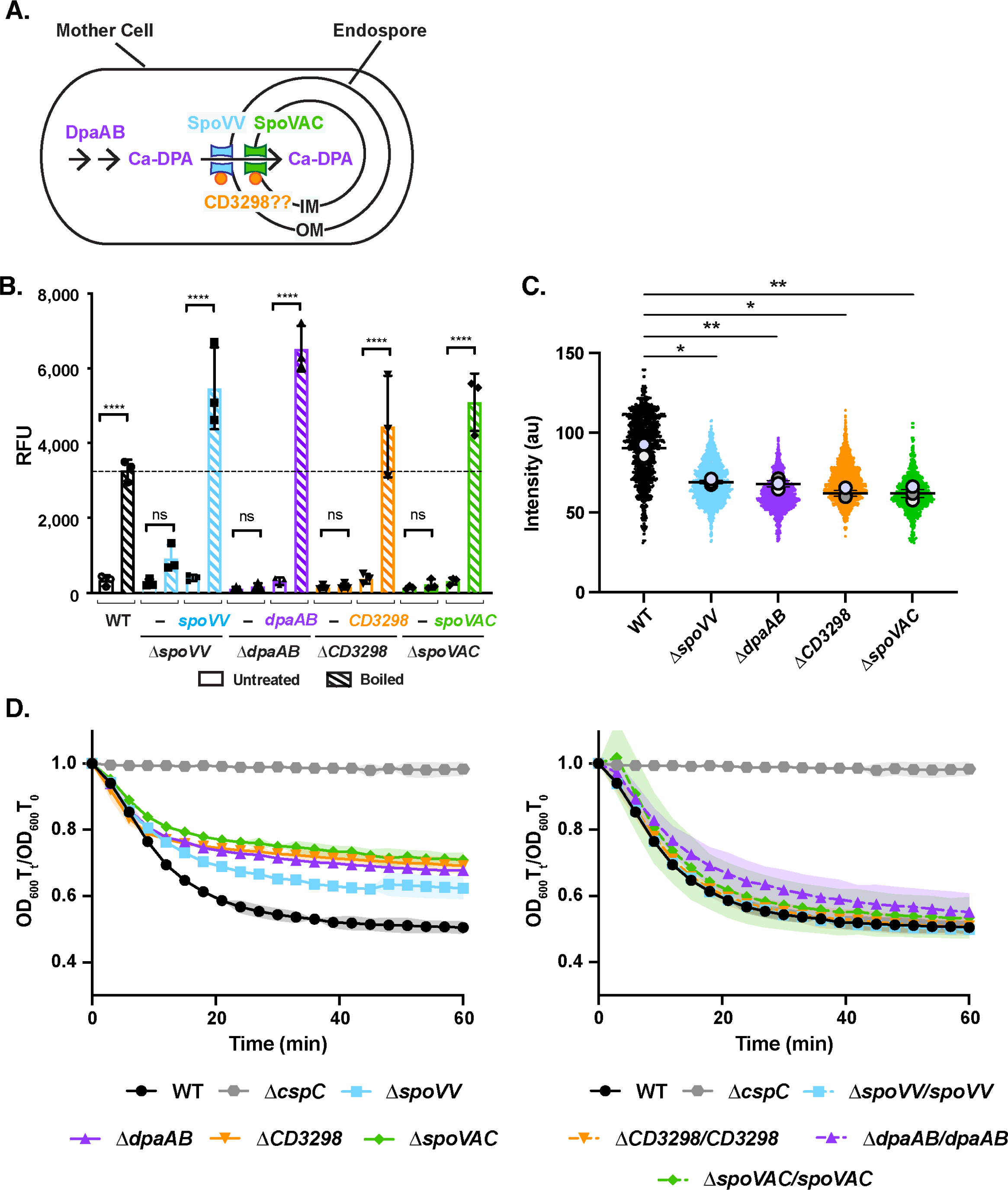
Mutants with defects in storing CaDPA in their core are more hydrated and less heat-resistant. (A) Schematic of CaDPA synthesis and transport during spore formation in *C. difficile*. DpaAB is the enzymatic complex that synthesizes DPA, which complexes with Ca^2+^ ions before it is transported across the outer (OM) and inner forespore membranes (IM). SpoVV is the putative CaDPA transporter in the OM (37), while SpoVAC is the CaDPA transporter located in the IM (43). CD3298 (orange circle) is a putative ATP-binding protein previously shown to be important for CaDPA transport in *C. difficile* (24), but it is unclear whether it associates with SpoVV or SpoVAC. (B) CaDPA levels measured in spores purified from WT, Δ*spoVV*, Δ*dpaAB*, Δ*cd3298*, Δ*spoVAC*, and their respective complementation strains. CaDPA levels were measured using terbium fluorescence after spores were boiled to release CaDPA or left untreated (RFU = relative fluorescent units). The average CaDPA measured on three independent spore preparations and associated standard deviation are shown. (C) SuperPlot comparison of CaDPA mutant spore intensity as measured by phase-contrast microscopy. The intensity of spores purified from the indicated strains was determined by phase-contrast microscopy using the MicrobeJ Fiji plug-in (44). A minimum of 650 spores was analyzed for each strain; the mean intensity for a given biological replicate is shown as large dots; the horizontal line represents the average of the mean intensity value for the three biological replicates. Statistical significance was determined using one-way ANOVA and Tukey’s test. (D) Optical density (OD_600_)-based analyses of spore germination over time in the indicated strains. Germination was induced in the presence of rich media (BHIS) and 1% (19 mM) taurocholate. The ratio of the OD_600_ of each strain at a given time point relative to the OD_600_ at time zero was plotted. The average of three independent experiments using three independent spore purifications is shown, and shaded error bars indicate the standard deviation for each time point measured. Statistical significance relative to WT was determined using a two-way ANOVA and Tukey’s test. **** p < 0.0001, *** p < 0.001, ** p < 0.01, * p < 0.05, n.s. = no statistical significance.

Consistent with prior work (24, 35), CaDPA was largely undetectable in Δ*dpaAB,* Δ*spoVAC* and Δ*cd3298* spores when spores were boiled to artificially release their CaDPA stores, and CaDPA was detected based on its terbium fluorescence (35, 45) (**Fig 1B**). Loss of SpoVV greatly reduced CaDPA levels, although Δ*spoVV* spores contained ∼5-fold higher levels of CaDPA than Δ*dpaAB* mutant spores. While this difference was not statistically significant, the elevated levels of CaDPA in Δ*spoVV* spores suggests that SpoVV may not be absolutely required for CaDPA transport into *C. difficile* spores. Importantly, complementation of the mutant strains with their respective genes from the *pyrE* locus under the control of their native promoters restored CaDPA levels within the spore core (**Fig 1B**), although Δ*dpaAB*/*dpaAB* spores had ∼two-fold higher CaDPA levels than WT 630Δ*erm* spores (35).

We next compared the spore purification efficiencies of the CaDPA mutant spores. We used a reduced Histodenz concentration because CaDPA mutant spores are less dense due to their greater hydration (35). Consistent with our prior analyses of JIR8094 strain *dpaAB* mutant spores (35), loss of DpaAB, SpoVAC, and CD3298 decreased spore purification efficiency by ∼two-fold (**Fig S1**), while loss of SpoVV did not affect the spore purification efficiency relative to WT. The complementation strains were purified at efficiencies comparable to WT, and ∼two-fold more Δ*dpaAB/dpaAB* spores were isolated relative to WT, in keeping with their two-fold higher levels of CaDPA relative to WT (**Fig S1**).

As a proxy for measuring core dehydration, we compared the intensity of our mutant spores during phase-contrast microscopy analyses because spores are more “phase-bright” when they are more dehydrated (35, 46, 47). By combining the MicrobeJ plug-in for ImageJ (44) with manual curation to measure the intensity of individual spores by phase-contrast microscopy, we found that CaDPA mutants were ∼25-30% less phase-bright than WT spores; Δ*CD3298* and Δ*spoVAC* spores were the least phase-bright on average (**Figs 1C, S1**).

Consistent with prior research indicating that core hydration is inversely correlated with heat resistance (35, 48, 49), CaDPA mutant spores were highly sensitive to heat treatment (**Fig S2**), whereas WT and complementation strain spores were resistant to 80°C heat treatment for 15 minutes. Δ*spoVV* spores were the most heat-resistant of the CaDPA mutant spores tested, since they maintained WT heat resistance up to 65°C, so the small amount of CaDPA present in Δ*spoVV* spores (**Fig 1B**) would appear to confer some wet heat resistance to these spores. Δ*CD3298* and Δ*spoVAC* spores were the least heat-resistant, with even a 15 min treatment at 50°C reducing their spore viability (**Fig S2**). The increased sensitivity of Δ*spoVAC* and Δ*CD3298* spores to heat relative to *dpaAB* mutant spores suggests that CD3298 may function alongside SpoVAC to transport molecules in addition to CaDPA during spore formation.

### CaDPA mutant spores appear to germinate less efficiently in optical density-based measurements

To assess the germination properties of CaDPA mutant spores, we used a well-established germination assay to monitor their decrease in optical density over time as spores degrade their cortex and hydrate their core. We previously used this assay to show that *dpaAB* and *spoVAC* mutant spores decrease in optical density less than WT spores because the mutant spores are more hydrated (35). Similar to these prior findings, the optical density of CaDPA-less (Δ*dpaAB*, Δ*spoVAC*, and Δ*CD3298*) spores decreased less than WT spores (35% vs. 50% decrease, respectively). Consistent with Δ*spoVV* spores containing some CaDPA (**Fig 1C**), the optical density decrease was Δ*spoVV* spores was intermediate between WT and CaDPA-less spores (**Fig 1D**). Δ*cspC* spores exhibited no change in OD_600_ because they lack the putative TA germinant receptor (20), while spores from the complementation strains exhibited WT-like reductions in OD_600_ (**Fig 1D**). Notably, although the relative OD_600_ decrease in CaDPA mutant spores was less than WT, their initial bulk germination rates (as defined as the average OD_600_ decrease over the first 6 min of germination) in rich media containing TA germinant were identical to WT irrespective of the TA concentration used. These data suggest that CaDPA mutants sense germinants and hydrolyze their cortex with efficiencies similar to WT in rich media containing TA germinant. These results are similar to what has been previously reported for the germination response of Δ*dpaAB* spores (35) and Δ*CD3298* (24) spores in rich media containing taurocholate.

### Spores lacking CaDPA do not require exogenous Ca^2+^ to germinate

Δ*CD3298* spores were previously reported to be incapable of germinating when germination is induced with TA germinant and glycine co-germinant alone (i.e. in the absence of endogenous or exogenous Ca^2+^, (24)), suggesting that Ca^2+^ is absolutely required for *C. difficile* spores to germinate. To rigorously test this proposal, we compared the germination responses of a panel of CaDPA mutant spores when germination is induced in the presence of TA germinant and either glycine vs. Ca^2+^ co-germinant. We initially used TA germinant and co-germinant concentrations that led to similar decreases in optical density as when we induced germination in rich media containing 1% (19 mM) TA for WT spores (**Fig 1D**). When CaDPA mutant spores were germinated in the presence of 1% (19 mM) TA and 10 mM glycine, they exhibited a similar loss in optical density (**Fig 2A**) as when they were induced to germinate in rich media (**Fig 1D**) or Ca^2+^ buffer containing TA germinant (**Fig 2B**). However, the initial rate at which CaDPA-less spores decrease in optical density was more gradual than for WT spores, especially for Δ*spoVAC* spores, indicating that CaDPA-less spores germinate more slowly as a population than WT spores in the presence of 1% (19 mM) TA and 10 mM glycine (**Fig 2A**). Regardless of the germinant and co-germinant concentrations used, i.e. rich media, TA + glycine, or TA + Ca^2+^, CaDPA mutant spores exhibited smaller decreases in optical density than WT spores (**Fig 1C**). Collectively, these results strongly suggest that Ca^2+^ is not absolutely required for *C. difficile* spore germination. However, CaDPA release appears to increase the average rate at which WT spores as a population germinate in response to TA germinant and glycine co-germinant based on bulk optical density-based germination analyses.

**Figure 2.**
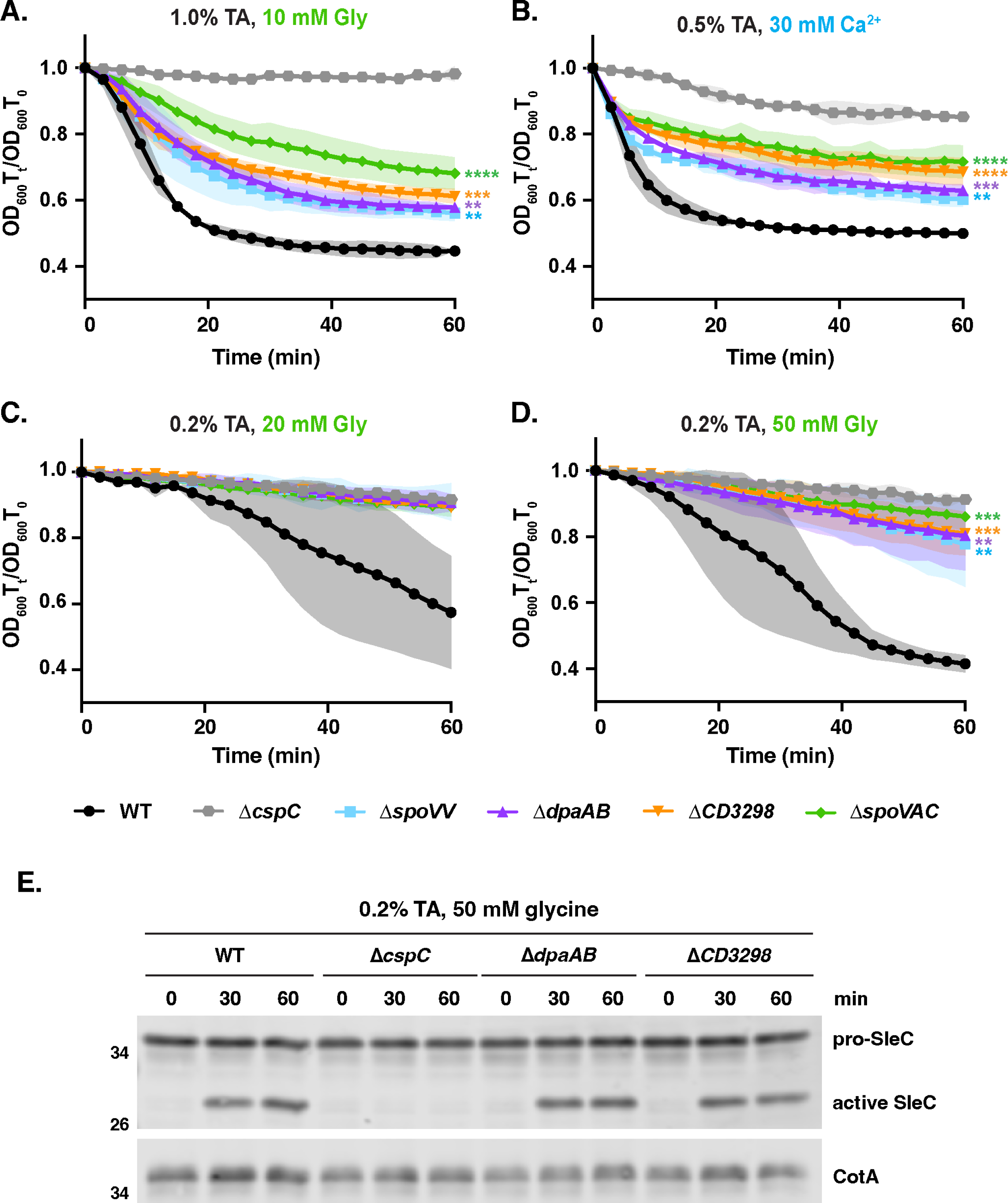
CaDPA mutant spores germinate in response to either Ca^2+^ or glycine co-germinants. Optical density (OD_600_)-based analyses of spore germination over time in the presence of the indicated concentrations of taurocholate and either glycine (A, C, D) or Ca^2+^ (B) co-germinants. The ratio of the OD_600_ of each strain at a given time point relative to at time zero was plotted. The average of three independent experiments (conducted on three independent spore purifications) is shown, and shaded error bars indicate the standard deviation for each time point measured. Statistical significance relative to WT was determined using a two-way ANOVA and Tukey’s test. **** p < 0.0001, *** p < 0.001, ** p < 0.01. (E) Western blot analyses of SleC activation following 0.2% (3.8 mM) TA and 50 mM glycine exposure in the indicated strains. The pro-SleC zymogen is proteolytically activated by the CspB protease in response to TA germinant and co-germinant signals (27, 28). CotA was used as a loading control (50).

Since these results contrasted with those of Kochan *et al.*, who failed to observe Δ*CD3298* spore germination in the presence of TA and glycine co-germinant alone in OD_600_-based assays, we repeated these analyses using the same conditions employed by these authors (24). Accordingly, we lowered the TA concentration to 0.2% (3.8 mM) and increased the glycine concentrations to 20 and 50 mM glycine. Under these conditions, WT spores germinated after a delay of ∼20 min in 20 mM glycine + 0.2% (3.8 mM) TA and ∼9 min in 50 mM glycine + 0.2% (3.8 mM) TA (**Figs 2C & D**). Although CaDPA mutant spores did not appear to germinate at the lower glycine co-germinant concentration (20 mM) with 0.2% (3.8 mM) TA relative to the Δ*cspC* control, their OD_600_ decreased by 10-15% after 60 min when glycine co-germinant concentrations were increased to 50 mM glycine with 0.2% (3.8 mM) TA, (**Fig 2D**, p < 0.01).

Since the increased hydration of CaDPA mutant spores complicates the interpretation of germination kinetics in the optical density-based assay, we analyzed germination using an optical density-independent method. To this end, we assessed the proteolytic activation of the cortex-degrading enzyme, SleC, by the CspB protease over time when germination was induced with TA germinant and glycine co-germinant in CaDPA-less mutant spores using western blotting. Under these conditions, the Δ*dpaAB* and Δ*CD3298* spores cleaved pro-SleC at levels indistinguishable from WT spores and with similar kinetics (**Fig 2E**), confirming that internal Ca^2+^ release is not required for *C. difficile* spores to initiate germination in the presence of TA and glycine co-germinant alone. Notably, Kochan *et al*. also observed some pro-SleC cleavage in Δ*cd3298* spores germinated in the presence of TA and glycine alone (24), highlighting how the optical density-based germination assay has reduced sensitivity in detecting germination in CaDPA mutant spores. Taken together, these data further support the conclusion that Ca^2+^ is not absolutely required to induce *C. difficile* spore germination as a co-germinant.

### Spores lacking CaDPA require higher levels of glycine co-germinant to germinate

Since CaDPA mutant spores appeared to germinate more slowly than WT spores in the presence of low levels of glycine co-germinant + TA germinant in the population-based optical density assays (**Figs 2A-B**), CaDPA release during *C. difficile* spore germination may serve to enhance TA + amino acid-induced spore germination. To directly test this possibility, we compared the bulk germination rates of CaDPA mutant spores in the presence of a fixed amount of TA germinant as we titrated either glycine or Ca^2+^ co-germinant. In the presence of increasing concentrations of Ca^2+^ and a fixed TA concentration (0.5%, 9.5 mM), the germination rate of WT spores increased steadily until it reached a maximum average rate at 7.5 mM Ca^2+^ (**Figs 3A-B**). Germination rate was defined as the average change in OD_600_ in the first 6 min of the assay. CaDPA mutant spores appeared to germinate at similar rates as WT spores at lower Ca^2+^ concentrations; however, at 7.5 mM Ca^2+^ and above, CaDPA mutant spores appeared to germinate more slowly than WT spores (**Fig 3B**). This apparent reduction in germination rate is likely due to the increased hydration of CaDPA mutant spores decreasing the dynamic range of the optical density-based germination assay (**Fig 3A**). Thus, CaDPA mutants would appear to exhibit WT responses to Ca^2+^ co-germinant in the presence of TA germinant.

**Figure 3.**
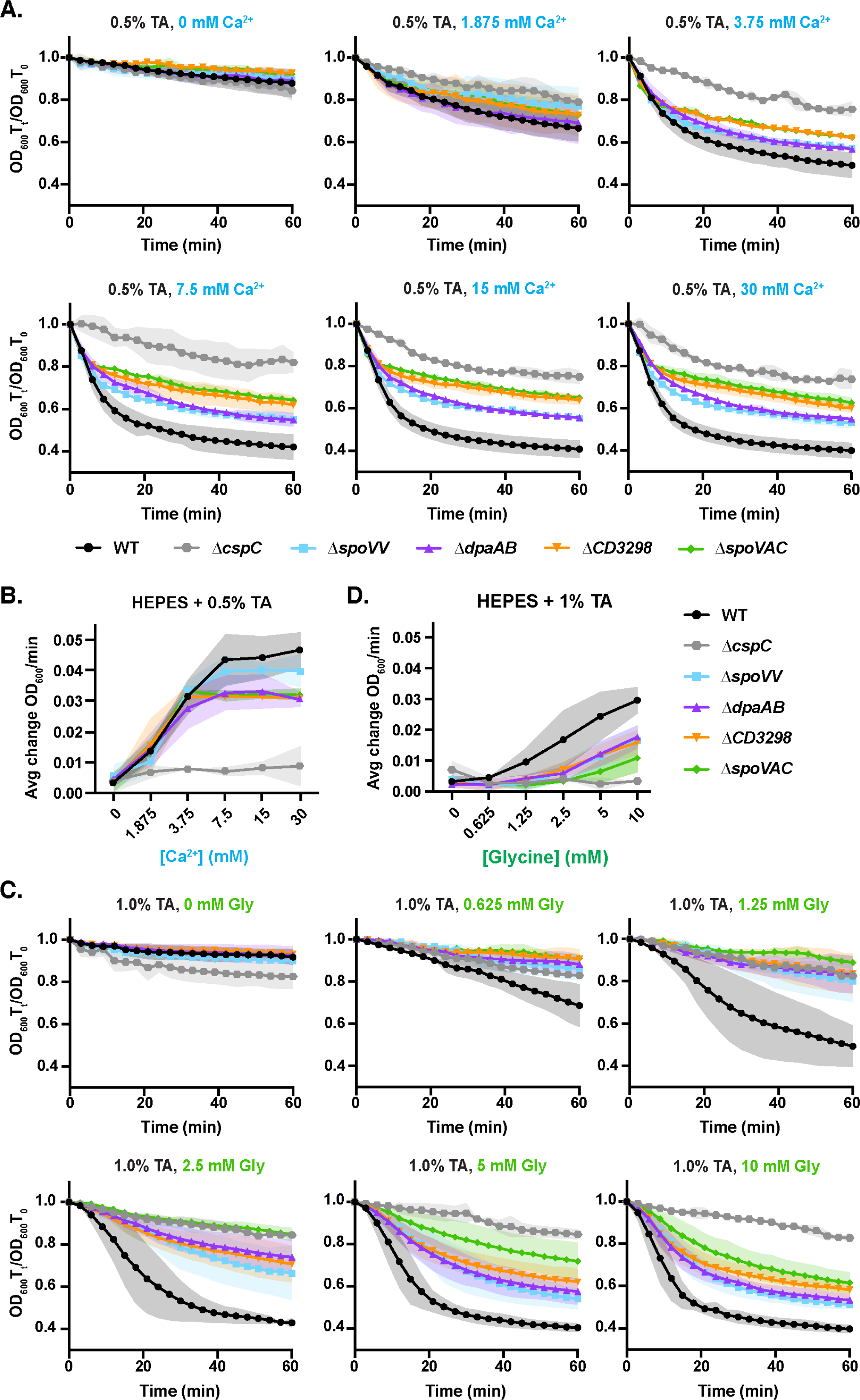
Sensitivity of CaDPA mutant spores to Ca^2+^ co-germinant. (A) Optical density (OD_600_)-based analyses of spore germination over time in the presence of the indicated concentrations of taurocholate and Ca^2+^. The ratio of the OD_600_ of each strain at a given time point relative to at time zero was plotted. (B) Germination rate as a function of Ca^2+^ co-germinant concentration. Germination rate was defined as the average change in OD_600_ in the first 6 min of the assay. (C) Optical density (OD_600_)-based analyses of spore germination over time in the presence of the indicated concentrations of taurocholate and glycine. The average of three independent experiments (conducted on three independent spore purifications) is shown, and shaded error bars indicate the standard deviation for each time point measured. (D) Germination rate as a function of glycine co-germinant concentration. Germination rate was defined as the average change in OD_600_ in the first 15 min of the assay. For all the data in this figure, the average of three independent experiments conducted on three independent spore purifications is shown, and the shaded error bars indicate the standard deviation for each time point measured.

We next assessed how CaDPA-less spores germinate relative to WT spores when germination is induced in the presence of increasing glycine co-germinant and a fixed TA concentration (1%, 19 mM TA). This higher concentration of TA was needed to reliably detect germination during the 60 min analysis time frame when germination was induced with TA + glycine co-germinant alone. Both WT and CaDPA-less spores exhibited a lag-phase to their germination, producing a sigmoidal vs. exponential decrease in OD_600_ (**Fig 3C**). Due to this lag, we expanded the time frame for which we calculated the average germination rate from 0 - 15 minutes rather than 0 - 6 minutes. Even when germination was induced with high glycine (10 mM) and 1% (19 mM) TA, CaDPA mutant spores appeared to germinate more slowly relative to WT (**Fig 3D**). The apparent germination rate of CaDPA mutant spores was decreased relative to WT spores at all glycine co-germinant concentrations tested (**Fig 3D**). Furthermore, CaDPA mutant spores required concentrations of 2.5 mM glycine to induce appreciable germination at 60 min in the optical density-based assay, whereas WT spore germination was detectable at 4-fold lower concentrations of glycine (0.625 mM) (**Figs 3C-D**). Taken together, our data indicate that CaDPA mutant spores are less sensitive to glycine co-germinant than WT spores. In addition, WT spores appear to germinate more slowly as a population when germination is induced in the presence of TA + glycine co-germinant compared to TA + Ca^2+^ co-germinant (**Fig 3**).

### Development of an automated image analysis pipeline for quantifying spore germination using time-lapse microscopy

Although the bulk germination analyses strongly suggested that internal Ca^2+^ can potentiate WT spore germination, they also underscored the challenges associated with measuring the germination kinetics of CaDPA mutant spores because the dynamic range of optical density-based germination assays is reduced in CaDPA mutant spores because of their hydrated nature. Notably, our western blot analyses of the proteolytic activation of SleC over time (**Fig 2E**) suggested that we may be missing germination events during the time frame of the OD-based assays since these bulk germination assays cannot detect if a small fraction of spores germinate within a population (< 5%, (51)). Furthermore, the assay can only be run for ∼60-90 min in a plate reader because spores start to settle over time. Indeed, the OD_600_ of Δ*cspC* spores decreases during this assay over time even though they are defective in inducing germination, as evidenced by their constant phase-bright intensity over time ((30), **Fig 4**). It is also difficult to distinguish between spores that initiate germination more slowly vs. heterogeneously using bulk optical density-based assays. Since directly monitoring the germination of individual spores using time-lapse microscopy would overcome many of these limitations, we developed an automated time-lapse microscopy germination assay for analyzing germination kinetics at the single-spore level (**Fig 4A**). When observed under phase-contrast microscopy, “phase-bright” dormant spores become “phase-dark” during germination as the cortex is degraded and the spore core becomes hydrated. To increase the number of spores that we could analyze with this microscopy-based assay, we developed an automated image analysis pipeline (PySpore) to analyze the germination of hundreds of individual spores in a given field-of-view. The PySpore program allows us to determine the time to germination (T_G_) and the germination rates of individual spores, with T_G_ being defined as when spores first start to decrease in phase-contrast intensity due to spore germination, and germination rate being defined as the maximum absolute rate of change of a 1-D smoothing spline fitted to the mean intensity of a given spore over the course of the assay. The maximum rate was calculated from the first derivative of the spline which was evaluated at each timepoint (**Fig S3**).

**Figure 4.**
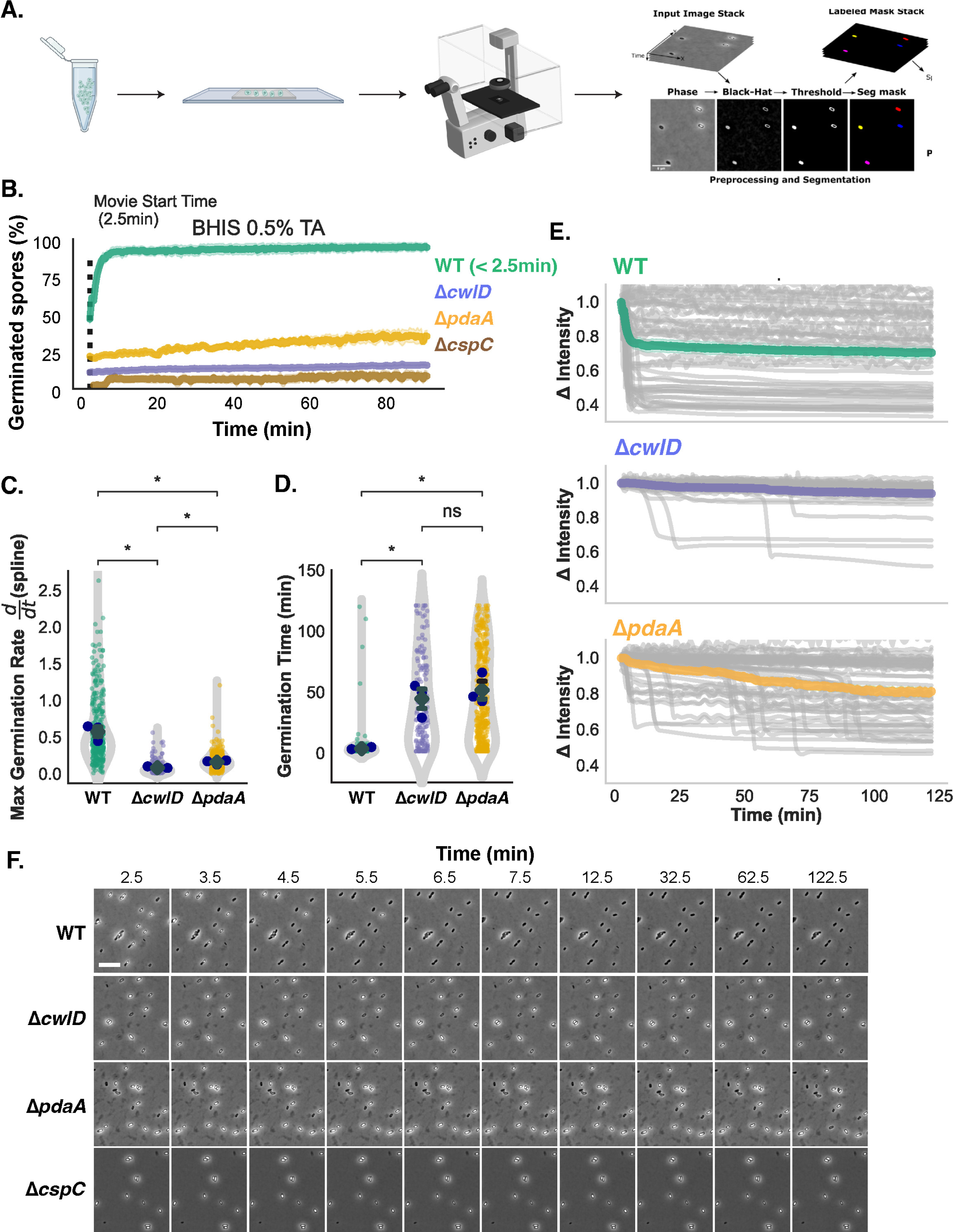
PySpore Automated image analysis pipeline for quantifying single-spore germination kinetics using time-lapse microscopy analyses. (A) Schematic of PySpore analyses of spore germination. Images from time-lapse microscopy are processed using a black-hat transform to identify spores based by enhancing their phase-dark outline. After spores are segmented, a simple single-particle tracking scheme allows them to be tracked throughout the course of the movie, and their intensities can be plotted over time. (B) Percent of germinated WT, Δ*cwlD*, Δ*pdaA*, and *ΔcspC* spores detected by the PySpore program for the indicated strains when spotted onto BHIS agarose pads containing 0.5%, 9.5 mM taurocholate. Spores whose intensity was below a specific threshold that did not change over time are included in the percent germinated value. Quantification of (C) time to germination and (D) maximum germination rate from PySpore analyses of the single-spore germination traces. (E) Single-spore germination traces of the change in relative intensity of WT, Δ*cwlD*, Δ*pdaA* spores. The ratio of the intensity at a given time point relative to the starting intensity is plotted. Statistical significance was determined via the Kruskal-Wallis test *p < 0.05. Blue points represent the median of each biological replicate, error bars show the mean (diamond) and standard error. (F) Filmstrip of time-lapse microscopy analyses of the indicated spores spotted on BHIS + 0.5%, 9.5 mM TA agarose pads. Parts of this figure were created with BioRender.

While single-spore germination assays have been developed previously (52, 53), developing *automated* analyses of individual germinating spores has been challenging for several reasons. WT dormant spores vary in their “phase-contrast” intensity, while the intensity (refractive index of the spore core) of germinating spores decreases over time as the spores become more hydrated (53)). As a result, it is difficult to set a universal threshold that correctly identifies all dormant spores at the start of a movie (**Fig 1C**). For example, when we tried to apply SporeTracker X, a previously developed automated image analysis program developed for *B. subtilis* spores (52), only ∼40% of *C. difficile* spores in our movie were correctly identified in the first frame (**Fig S4**).

To overcome the thresholding issue, we reasoned that the fixed intensity of the phase-dark spore coat would allow us to segment and track spores from frame-to-frame even as the intensity of their core as measured by phase-contrast microscopy changes over time during germination (**Fig 4A**). By segmenting spores based on their dark outline, we were able to identify >90% of spores in a given frame, improving the fidelity of spore segmentation by two-fold over SporeTrackerX (**Fig S4**, (52)).

To validate our PySpore analysis method, we analyzed the germination of *C. difficile* mutant spores with defects in cortex hydrolysis. Δ*cwlD* and Δ*pdaA* cortex modification mutants degrade their cortex layer more slowly because they lack muramic-∂-lactam, a spore-specific peptidoglycan modification that is recognized by the main cortex lytic enzyme, SleC (51, 54). In the optical density-based bulk germination assay, Δ*cwlD* spores do not appear to germinate because their OD_600_ stays largely constant, while the OD_600_ of Δ*pdaA* spores decreases slightly (∼10% after 3 hrs). When we analyzed the germination properties of these mutant spores on agarose pads containing BHIS and 0.5% (9.5 mM) TA relative to WT spores using time-lapse microscopy (**Fig 4B**), we found that a small percentage (∼5%) of Δ*cwlD* spores germinated during the 2 hr assay period (**Fig 4C**), with germinating spores being defined as those that decrease in intensity by >20%. In contrast, ∼40% of Δ*pdaA* spores germinated over the 2 hr assay albeit with considerably delayed kinetics relative to WT spores. Indeed, WT spores germinated rapidly and uniformly under these conditions, with most WT spores completing germination within ∼5 min of being exposed to germinant. In contrast, the median “time to germination” for Δ*pdaA* and Δ*cwlD* spores was ∼50 min (**Fig 4D**). Furthermore, these mutant spores germinated more slowly at the single-spore level, with the median germination rate for a given spore being ∼3-fold slower than the rate measured for WT spores (**Fig 4E**). It should be noted that many WT spores had already completed germination within the 2.5 min required to set up a movie (**Figs 4C, 4F**), so the median germination time shown for WT spores is an underestimate. Spores that had already germinated were identified by setting an intensity threshold to identify phase-dark spores whose intensity did not change over the assay length but had clearly completed germination. Regardless, the slower germination rate and delays in time to initiation of Δ*cwlD* and Δ*pdaA* spores are consistent with their cortex modification defects, which decreases the efficiency with which the SleC cortex lytic enzyme degrades the unmodified cortex of these mutant spores (51, 54). Since germinant sensing is unaffected in Δ*cwlD* and Δ*pdaA* spores, the delays in the “time to germination” and the slower germination rates measured for these mutant spores likely reflect how both time-lapse microscopy and optical density-based germination assays depend on the cortex being degraded and the core becoming hydrated in order for changes in spore intensity or optical density to be detected (51, 54). As a result, the “time to germination” measurement reflects the initiation of germination (i.e. germinant and co-germinant signal transduction), degradation of the cortex, and hydration of the core. Regardless, these observations indicate that our time-lapse, single-spore germination assay yields data consistent with prior optical density-based germination analyses of Δ*cwlD* and Δ*pdaA* spores (51). Furthermore, the single-spore analyses (**Fig 4F**) provide additional insight into the germination properties of these cortex modification mutant spores by sensitively revealing their heterogeneous germination, delayed time to germination, and reduced germination rate.

### Single-spore time-lapse germination analyses of Δ*dpaAB* spores reveal differences in their germination responses to glycine vs. Ca^2+^ co-germinants

Having established the utility of our single-spore germination assay, we next used it to analyze the germination of CaDPA-less spores in the presence of TA + glycine alone (i.e. the absence of added Ca^2+^). WT and Δ*dpaAB* spores were induced to germinate in the presence of 0.2% (3.8 mM) TA with 10 mM glycine. The TA concentration was lowered to 0.2% (3.8 mM) to capture most of the phase-bright spores transitioning to phase-dark spores within the 2.5 min needed to set up the time-lapse microscopy movies. In the presence of 0.2% (3.8 mM) TA and 10 mM glycine, approximately half as many Δ*dpaAB* spores germinated during the assay as WT spores (**Fig 5A**), and their germination rate, as defined by the decrease in phase intensity over time, was slower relative to WT (**Fig 5B**).

**Figure 5.**
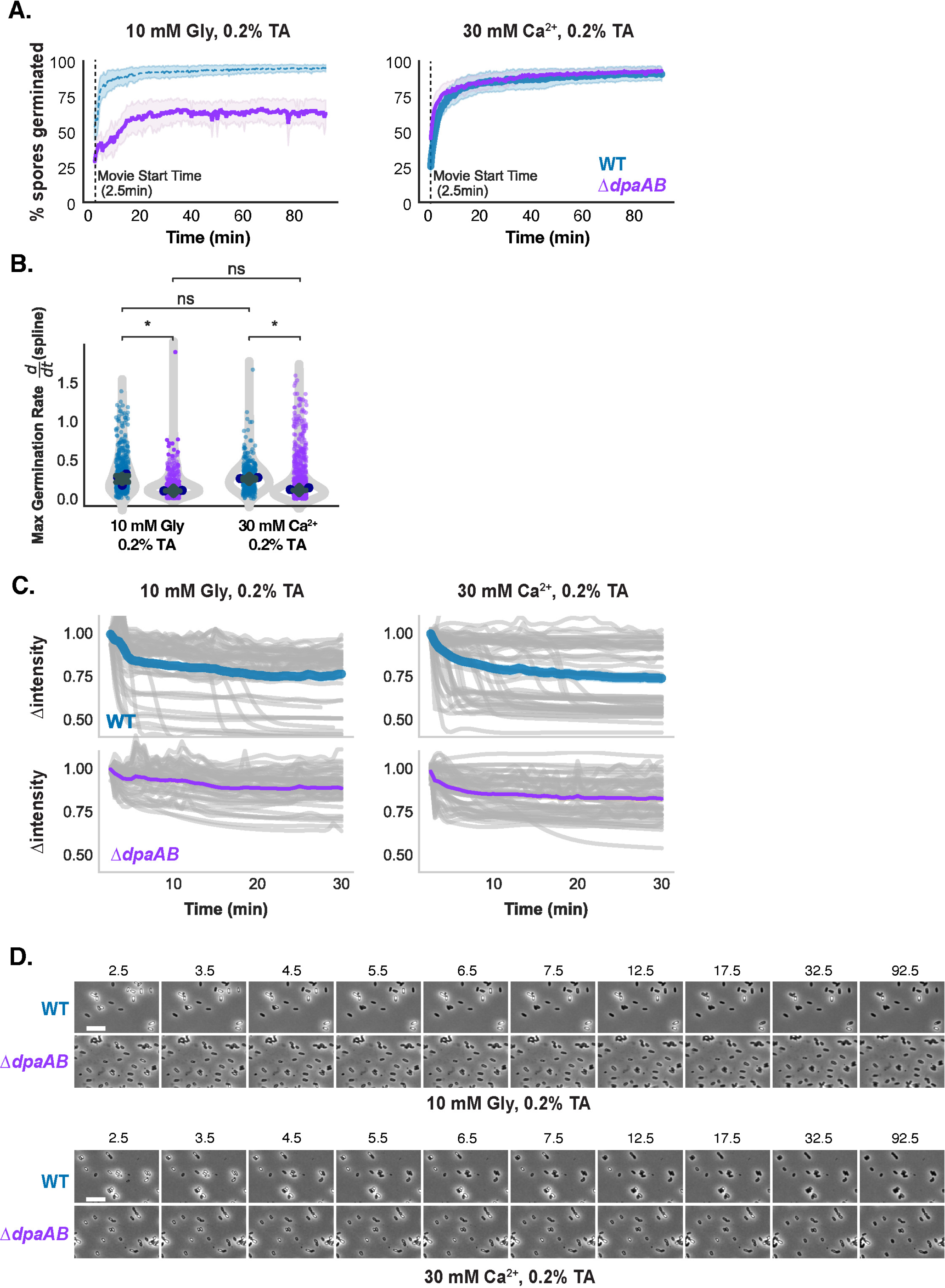
Single-spore analysis of the individual roles of Ca^2+^ and Gly during germination. (A) Kinetic analysis of percent germination of WT and Δ*dpaAB* spores in HEPES-NaCl buffer containing 0.2% (3.8 mM) TA and 10 mM glycine or 30 mM CaCl_2_. (B) Comparison of maximal germination rate for WT and Δ*dpaAB* with either glycine or calcium alone. (C) Single germination traces showing the change in intensity over time under each condition. (D) Filmstrips of WT and Δ*dpaAB* in each buffer condition (scale bar = 5 μm). Statistical significance was determined via the Kruskal-Wallis test * p < 0.05. Blue points represent the median of each biological replicate, error bars show the mean of all biological replicates (grey diamond) and standard error.

Interestingly, the germination rates measured for WT spores were more heterogenous relative to Δ*dpaAB* spores, perhaps because CaDPA released from other WT spores can promote the germination of neighboring spores. Consistent with this hypothesis, no difference in the proportion of germinating Δ*dpaAB* and WT spores was observed over time using the single-spore assay when germination was induced with 0.2% (3.8 mM) TA and 30 mM Ca^2+^ (**Fig 5A**). Despite these similarities, individual Δ*dpaAB* spores still germinated more slowly than WT spores, similar to our observations with TA + glycine-induced germination (**Figs 5B-C**). Again, this slower apparent germination rate is likely due to the greater hydration of CaDPA mutant spores, which reduces the concentration gradient between the spore core relative to the surrounding media such that *ΔdpaAB* spores likely do not hydrate as quickly as WT spores. Importantly, the aggregated single-spore germination data (**Fig 5C**) was similar to bulk optical density-based germination analyses (**Fig S5**).

While these observations highlight the complexities in measuring the germination of CaDPA mutant spores using phase-contrast microscopy, the single-spore germination data (i) clearly demonstrate that these mutant spores can germinate in the presence of TA + glycine alone, i.e. in the absence of added Ca^2+^ and (ii) suggest that internal Ca^2+^ release can potentiate the germination of other spores within the population.

### Ca^2+^ released from the spore core during germination is a “public good” that can be sensed by other spores

To directly test the hypothesis that CaDPA released by spores can serve as a “public good” to promote the germination of other spores in the population, we compared the ability of germination “exudates” released from germinating WT and Δ*dpaAB* spores to induce the germination of “naïve” WT spores (i.e. not previously exposed to germinant) in our time-lapse microscopy assay. WT and Δ*dpaAB* spores were induced to germinate in the presence of low glycine (2.5 mM) and TA (0.5%, 9.5 mM) concentrations for 16 hrs. This extended incubation time was necessary to allow the population of Δ*dpaAB* spores to fully germinate, since they complete germination more slowly than WT cells under these conditions (**Fig 6A**). During this incubation time, WT spores release CaDPA into the supernatant fraction, while Δ*dpaAB* do not. We reasoned that WT germination exudates, which contain CaDPA released by these spores, would promote the germination of “naïve” WT spores more readily when incorporated into agarose pads than germination exudates of Δ*dpaAB* spores, which do not contain released CaDPA (**Fig 6A**). Indeed, when “naïve” WT spores were spotted onto agarose pads made from WT germination exudates, a greater proportion of these spores induced germination relative to spores spotted onto pads made of mock-treated buffer, which contains the 2.5 mM glycine and 0.5% (9.5 mM) TA originally used to prepare the WT and Δ*dpaAB* germination exudates. Similarly, fewer WT spores germinated when spotted onto pads made with germination exudate prepared from Δ*dpaAB* spores, which lacks CaDPA in the exudate fraction (**Fig 6B**). These findings strongly suggest that CaDPA released into WT germination exudates potentiates the germination of other spores in the population.

**Figure 6.**
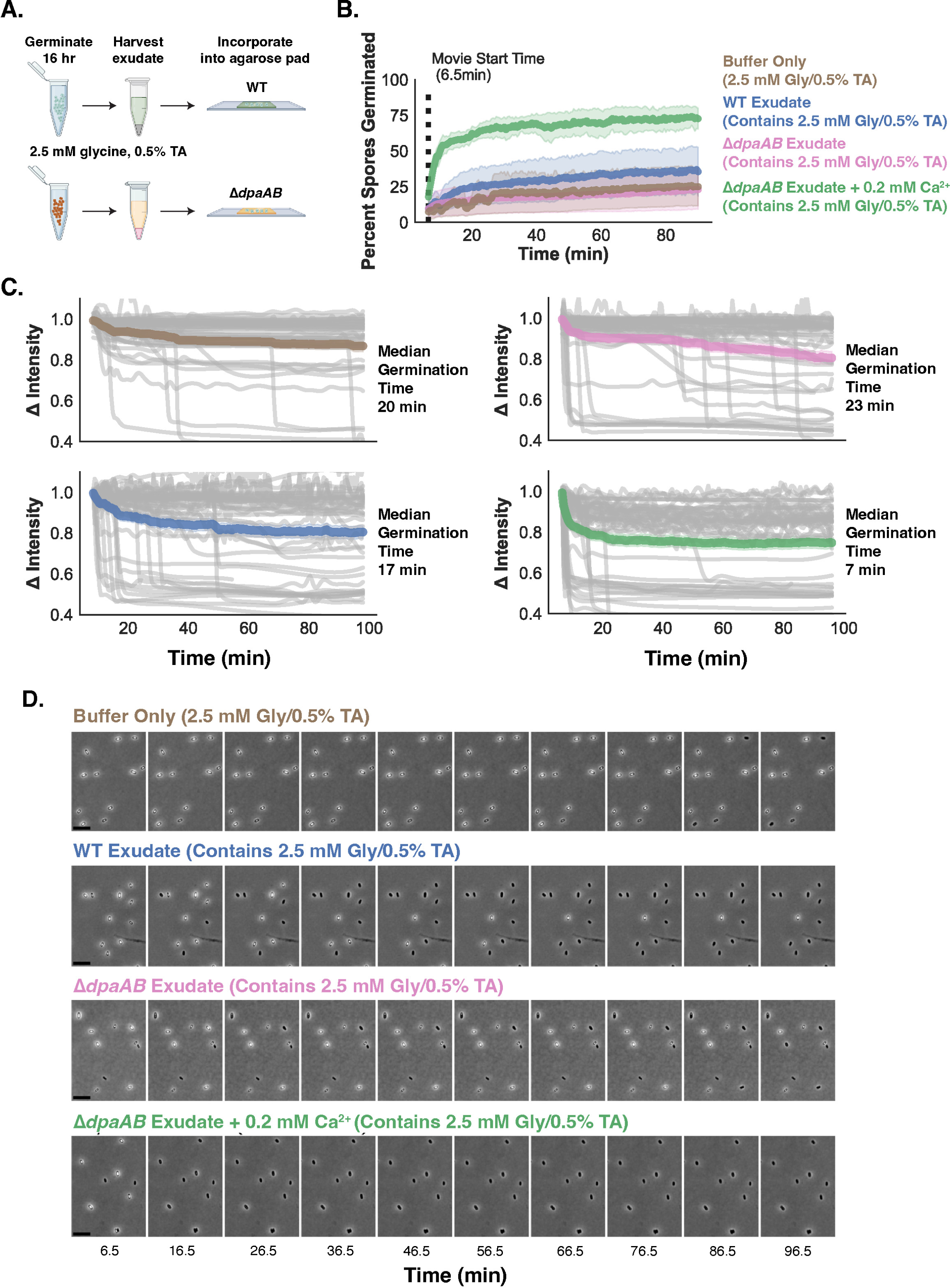
Ca^2+^ released during WT spore germination can serve as a “public good” to potentiate the germination of neighboring spores. (A) Schematic of germination exudate preparation. Phase-contrast microscopy images highlight the delayed germination kinetics of Δ*dpaAB* spores in the presence of low glycine and TA. After germinating WT and Δ*dpaAB* spores in 2.5 mM glycine and 0.5% (9.5 mM) TA, germination “exudates” were harvested by pelleting spores and removing the supernatant fraction. This fraction was incorporated into agarose pads, and the ability of CaDPA released into the WT germination exudate fraction to potentiate germination in “naïve” WT spores is assessed. (B) Percent of germinated spores detected by the PySpore program for the indicated strains when spotted onto BHIS agarose pads containing 0.5% (9.5 mM) taurocholate. Spores whose intensity was below a specific threshold that did not change over time are included in the percent germinated value. (C) Single spore traces showing the change in intensity over time for WT spores in the presence of each harvested exudate. (D) Filmstrips of germination for WT spores germinating in the presence of harvested exudates.

To more directly link the ability of WT germination exudates to promote germination to the CaDPA released into these extracts, we supplemented Δ*dpaAB* germination exudates with Ca^2+^ as a proxy for CaDPA (Ca^2+^ is in a one-to-one complex with DPA in the core). To match the Ca^2+^ levels in WT germination exudates, we first measured the DPA concentration in WT germination exudates using terbium chloride. These analyses indicated that supplementing Δ*dpaAB* germination exudates with 0.2 mM calcium chloride should mimic the concentration of Ca^2+^ in WT germination exudates. When 0.2 mM CaCl_2_ was added to Δ*dpaAB* germination exudates, the supplemented agarose pads rapidly induced germination in the population of WT spores. Since sub-millimolar levels of Ca^2+^ were able to reduce median germination times by ∼3-fold, these data highlight how Ca^2+^ and glycine co-germinants can synergize to induce WT spore germination, similar to a previous report by Kochan *et al*. (24). Taken together, our data indicate that internal Ca^2+^ released from germinating spores potentiates the germination of other spores within the population. Accordingly, CaDPA-less spores are less sensitive to glycine co-germinant because they do not release CaDPA and thus cannot potentiate the germination of other spores within the population.

### Ca^2+^ appears to be required for a step downstream of (co-)germinant signaling

Although we used multiple germination analyses to demonstrate that endogenous Ca^2+^ release is not necessary for germination stimulated in the presence of TA + glycine alone (Figs 2, 3, 5, and 6), Ca^2+^ may still be needed at low concentrations to regulate a factor required for a germination step downstream of germinant and co-germinant signaling. Kochan *et al*. provided evidence that Ca^2+^ may be needed as a co-factor when they showed that EGTA at a concentration < 50 µM was sufficient to inhibit germination induced by 0.2% TA and 50 mM glycine (Fig 2A). Addition of excess Ca^2+^ (1 mM) was sufficient to counteract the effects of EGTA (Fig 2B, Kochan *et al*. 2017) and restore spore germination. Taken together, these analyses with EGTA suggest that Ca^2+^ is required at concentrations < 50 µM for germination for germination to proceed if Ca^2+^ serves as a co-factor.

In contrast, for Ca^2+^ to serve as the sole co-germinant source to promote TA-induced germination, Ca^2+^ must be supplied at 2-5 mM concentrations (i.e. 100-fold higher) to function as a co-germinant to promote germination at ∼50% maximal levels in the presence of 0.2-0.5% (3.8-9.5 mM) TA (**Fig 3B**, Kochan *et al*. 2017). However, it is also important to note that the Ca^2+^ and glycine co-germinants can synergize with TA germinant such that low µM levels of Ca^2+^ can stimulate germination in the presence of glycine and TA (Kochan *et al*. 2019, **Fig 6B** current manuscript). As a result, it can be challenging to distinguish between the ability of Ca^2+^ to potentiate germination as a co-germinant versus its requirement as a co-factor for germination to proceed.

To attempt to distinguish between these possibilities, we titrated the Ca^2+^ chelator, EGTA, to determine the minimum concentration needed to inhibit the germination of WT and Δ*dpaAB* spores. For these experiments, we induced germination using the same conditions used by Kochan *et al*., who previously showed that 50 µM EGTA potently inhibits WT spore germination induced by 0.2% (3.8 mM) TA and 50 mM glycine (Kochan *et al*. 2017). These analyses revealed that EGTA inhibited germination at concentrations between 2 and 10 µM EGTA for both WT and Δ*dpaAB* spores (newly added **Fig 7A**). Interestingly, at 10 µM EGTA, we observed that some WT spore preparations induced germination ∼70 min into the 90 min assay, suggesting that some WT spores are able to release CaDPA to sufficient levels to promote germination to function in the positive feedback loop described in Fig 6. Taken together, these analyses indicate that Ca^2+^ is needed at concentrations between 2-10 µM for germination to proceed under the conditions tested.

**Figure 7.**
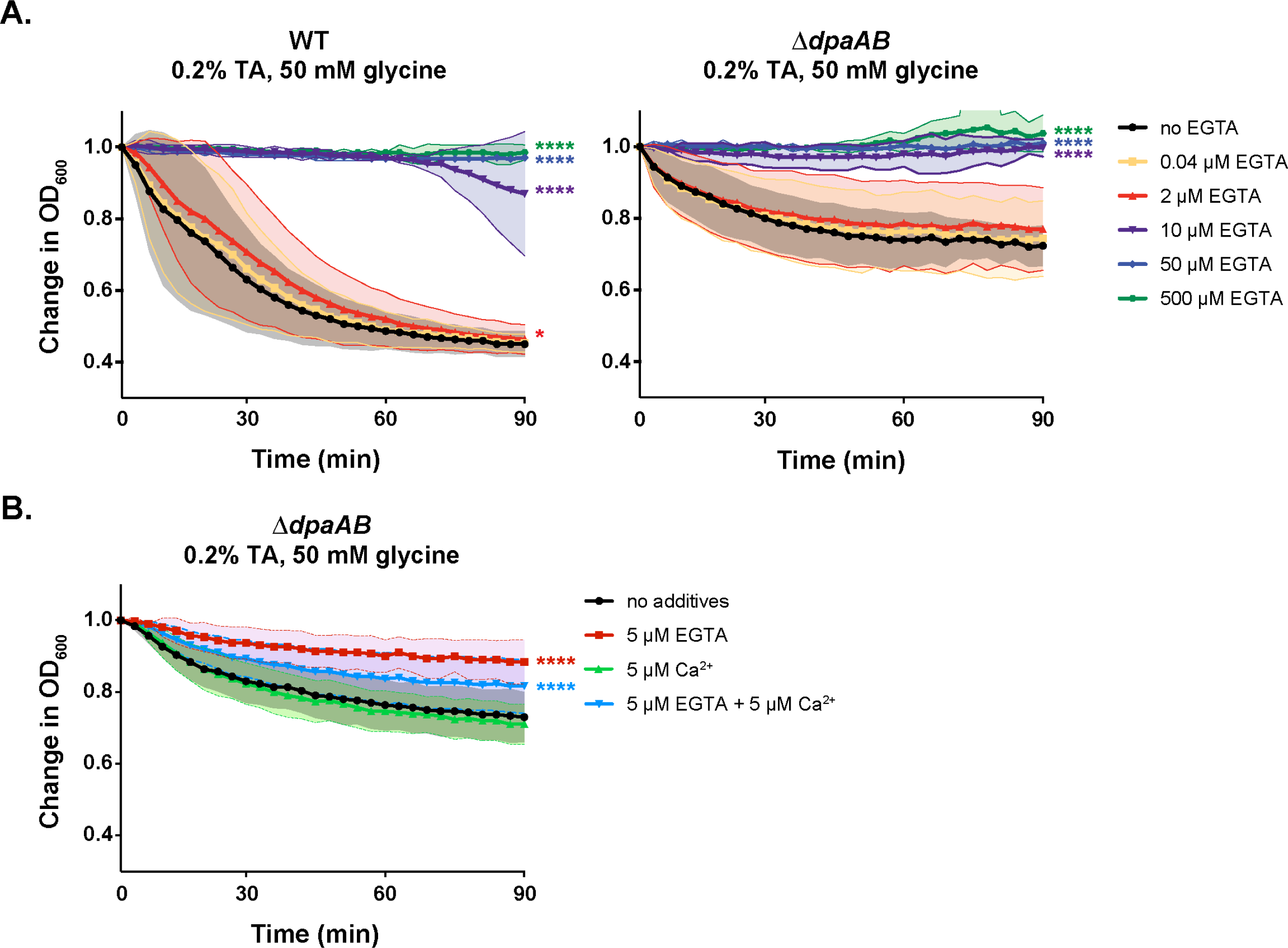
Micromolar concentrations of Ca^2+^ appear to be required for *C. difficile* germination to proceed. Optical density (OD_600_)-based analyses of WT and Δ*dpaAB* spore germination over time in the presence of 0.2% (3.8 mM) taurocholate and 50 mM glycine and the indicated concentrations of EGTA (A) and 5 µM EGTA and/or 5 µM Ca^2+^ (B). The ratio of the OD_600_ of each strain at a given time point relative to at time zero was plotted. The average of three independent experiments (conducted on three independent spore purifications) is shown, and shaded error bars indicate the standard deviation for each time point measured. Statistical significance relative to the untreated sample was determined using a two-way ANOVA and Tukey’s test.

To more directly assess whether Ca^2+^ functions as a co-germinant to stimulate germination versus as a co-factor required for germination to proceed when induced by TA + glycine, we compared the effect of chelating Ca^2+^ using 5 µM EGTA versus adding Ca^2+^ as a co-germinant. Since we observed variation in the levels of inhibition by EGTA at 5 µM in WT spores (**Fig S6**) (presumably due to the variable of endogenous Ca^2+^ release in these analyses participating in a feedforward loop, Fig. 7A), we determined the effect of 5 µM EGTA vs. Ca^2+^ on Δ*dpaAB* spores only. In the presence of 0.2% (3.8 mM) TA and 50 mM glycine, 5 µM EGTA largely inhibited germination, while 5 µM CaCl_2_ supplementation did not promote germination above the TA + glycine alone condition (**Fig 7B**). When both 5 µM EGTA and 5 µM CaCl_2_ were added to Δ*dpaAB* spores induced to germinate in the presence of 0.2% (3.8 mM) TA and 50 mM glycine, CaCl_2_ was able to partially counteract the inhibitory effect of EGTA (**Fig 7B**). Since low µM concentrations of Ca^2+^ were insufficient to synergize with the glycine co-germinant present in the assay, whereas equivalent concentrations of the Ca^2+^ chelator EGTA were able to inhibit germination under these conditions, these results are consistent with Ca^2+^ being required for a germination step downstream of germinant and co-germinant signaling. Thus, it is likely that Ca^2+^ is required as a co-factor for a protein critical for germination to proceed.

## Discussion

Unlike previously studied spore-formers, the nosocomial pathogen *C. difficile* requires multiple environmental signals to induce germination (11, 13, 18). While *C. difficile*’s use of bile acids as a germinant signal has been well-established (23), the precise requirement for amino acid and divalent cation co-germinants during *C. difficile* germinant sensing has been a matter of debate, with some groups proposing that divalent cations, namely Ca^2+^, are essential for *C. difficile* germination (24). In this study, we used a panel of CaDPA mutants and several independent germination assays (optical density, single-spore, time-lapse microscopy, and western blotting, **Figs 2, 4, 5**) to show that Ca^2+^ is not absolutely required as a co-germinant signal for *C. difficile* spore germination. Specifically, we demonstrate that *C. difficile* mutants that lack internal Ca^2+^ stores in the form of CaDPA can still germinate when exposed to bile acid and amino acid co-germinants alone. However, under these conditions, CaDPA mutant spores require higher concentrations of amino acid co-germinants and longer incubation times (**Figs 2,5**). While our analyses indicate that Ca^2+^ is not required as a co-germinant signal to induce germination stimulated by TA + glycine alone, our analyses of germination induced by TA + glycine in the presence of EGTA and/or exogenous Ca^2+^ suggest that trace levels of Ca^2+^ are required for a germination step downstream of germinant sensing and signaling. Thus, Ca^2+^ is likely needed as a co-factor for an enzyme required during germination (but not germinant or co-germinant sensing).

Notably, these data indicate that *either* class of co-germinant can stimulate *C. difficile* spore germination when combined with TA germinant, our findings strongly suggest that these two distinct classes of co-germinants are sensed by distinct mechanisms in *C. difficile*. This conclusion is supported by our prior genetic data, which identified specific substitutions in the CspC bile acid germinant receptor that sensitize *C. difficile* spores to specific classes of co-germinant (30). While the molecular basis for distinguishing between these distinct co-germinant classes is unclear, the CspA pseudoprotease clearly plays an important role in regulating co-germinant sensing given that certain CspA mutations bypass the need for either class of co-germinant signals (28). Since our data indicate that parallel pathways mediate co-germinant sensing, CspA may function downstream of the actual co-germinant sensor rather than as the direct sensor of both classes of co-germinants. Distinguishing between these different models awaits future biochemical identification of the direct sensors of amino acids and Ca^2+^, respectively. Regardless, our data suggest that distinct regions of the same receptor or two distinct receptors are involved in sensing amino acid and Ca^2+^ co-germinant signals, respectively.

Notably, rigorously assessing the requirement for Ca^2+^ as a co-germinant signal during *C. difficile* spore germination required that we optimize a method for directly monitoring spore germination at the single-cell level (**Fig 3**). This method was necessary because the lack of CaDPA in these mutant spores made it difficult to use traditional optical density-based assays to accurately monitor their germination over time. By coupling time-lapse microscopy analyses of individual spores to our custom image analysis pipeline (**Fig 4**), we were able to demonstrate that CaDPA-less spores still germinate when germination is induced by TA + glycine alone, i.e. in the absence of added Ca^2+^, although a smaller fraction of Δ*dpaAB* spores germinated under these conditions relative to WT or to Δ*dpaAB* spores germinated with taurocholate and calcium (**Fig 4**), highlighting how CaDPA release can potentiate germination of other spores within the population. Indeed, our germination exudate experiments allowed us to demonstrate that CaDPA released during *C. difficile* spore germination forms a feedforward loop that potentiates the germination of neighboring spores when germination is induced by TA and amino acid germinants alone (**Fig 5**). While the physiological significance of this feedforward loop remains unclear, it is possible that local changes in Ca^2+^ within the gut could promote *C. difficile* spore germination, especially glycine was recently shown to be a limiting factor for establishing infection (55). Indeed, it is possible that spores from other bacterial species could potentiate *C. difficile* spore germination, since CaDPA is released by all germinating spores. Unfortunately, testing whether CaDPA release by neighboring spores contributes to germination *in vivo* could be challenging given that CaDPA-less spores have decreased resistance properties and may not survive passage through the stomach.

Our analyses of several strains with CaDPA defects also provide new insight into how CaDPA is incorporated into *C. difficile* spores. Our data confirm that SpoVV regulates CaDPA transport in *C. difficile* likely by allowing CaDPA to cross the mother cell-derived outer forespore membrane as previously shown in *B. subtilis* (37). However, in contrast with *B. subtilis*, some CaDPA can still be transported into *C. difficile* spores in the absence of SpoVV, implying that there is an additional transporter that can partially substitute the loss of its function. In addition, our data suggest that CD3298, a putative ATP-binding protein, regulates SpoVAC function, since the resistance and germination phenotypes of Δ*cd3298* spores most closely resemble those of Δ*spoVAC* spores. Alternatively, CD3298 may work in concert with SpoVV and the unidentified alternative CaDPA transporter located in the mother cell-derived outer forespore membrane.

Taken together, our findings resolve an important question in our understanding of how *C. difficile* integrates signals from amino acid and Ca^2+^ co-germinants alongside bile acid germinants by demonstrating that either co-germinant class is sufficient to function as a co-germinant. They also provide a new quantitative method for monitoring germination kinetics over time, since our automated PySpore method allows the germination of hundreds to thousands of spores to be analyzed over time. Since this method is not specific to *C. difficile*, it can be applied to study mechanisms of germination initiation and germination heterogeneity in other spore-formers with greater accuracy than existing programs (**Fig S4**). The ease with which this program can be implemented by other investigators and its improved accuracy in identifying spores within a population will hopefully allow the method to be adopted more widely.

## Materials and Methods

### Bacterial strains and growth conditions

The parental strain for all *C. difficile* strains in the study was 630Δ*ermΔpyrE*. Allele coupled exchange (ACE), which allows for in-frame gene deletions and complementation in an ectopic locus, was used to generate all strains (56). *C. difficile* strains are listed in **Table S1** were grown on brain heart infusion (BHI) media supplemented with L-cysteine (0.1% w/v; 8.25 mM), taurocholate (TA, 0.1% w/v, 1.9 mM), thiamphenicol (5-15 µg/mL), kanamycin (50 µgmL) or cefoxitin (8 µg/mL), as needed. For constructing in-frame deletions using allele coupled exchange, *C. difficile* defined media (CDDM, (57)) was supplemented with 5-flouroorotic acid (5-FOA, 2 mg/mL) and uracil (5 µg/mL). Cultures were grown at 37°C under anaerobic conditions using a gas mixture containing 85% N_2_, 5% CO_2_ and 5-10% H_2_.

*Escherichia coli* DH5α strain was used for all plasmid construction and *E. coli* strain HB101/pRK24 was used for conjugating plasmids for ACE into *C. difficile*. All HB101 strains used in this study are listed in **Table S1**. E*. coli* strains were grown in Luria-Bertani broth (LB) at 37°C with shaking at 225 rpm. The media was supplemented with chloramphenicol (20 µg/mL) or ampicillin (50 µg/mL) as indicated.

### *E. coli* strain construction

Plasmid constructs were cloned into DH5α strains and sequenced confirmed using Genewiz Sanger sequencing. Benchling plasmid maps for all constructs and primers used in strain construction are included in **Table S2**. PCR products were cloned into pMTL-YN3 digested with AscI and Sbf1 or pMTL-YN1C digested with NotI and XhoI using Gibson assembly. Sequenced confirmed plasmids were then cloned into HB101 strains to conjugate into *C. difficile* strains.

### *C. difficile* strain construction

*C. difficile* in-frame deletions strains were generated with ACE by conjugating pMTL-YN3 plasmid constructs in HB101 into the 630Δ*erm*Δ*pryE* parental strain. Deletions were confirmed by diagnostic PCR and two individual clones were phenotypically characterized before restoring the *pyrE* locus. The *pyrE* locus was restored with ACE by conjugating the pMTL-YN1C plasmid into the deletion strain or through complementation with appropriate pMTL-YN1C plasmid containing the gene of interest. Two individual complementation clones were phenotypically characterized. All strains used for conjugations are included in Table S1.

### Spore purification

Strains were grown on 70:30 agar media for 2-3 days to induce sporulation as previously described (58). Cultures were harvested into ice-cold, sterile water. Spore samples were washed 5-6 times in ice-cold, sterile water and incubated overnight in sterile water at 4°C. After incubation, spore solutions were washed in ice-cold, sterile water and then treated with DNase I (New England Biolabs) for 60 mins at 37°C. Spores were then purified on a 20%/45% Histodenz (Sigma Aldrich) gradient. The 45% Histodenz was required to isolate the less dense, CaDPA-less spores (35). The resulting spores were washed 2 times in ice-cold, sterile water and assessed for purity using phase-contrast microscopy. Spore stocks were stored at 4°C in sterile water. OD_600_ of the spore stock was measured as a proxy for spore concentration. The average spore purification yield per agarose plate was determined from three independent spore purifications.

### Total DPA quantification using terbium fluorescence

The total DPA content of spores were measured using terbium fluorescence essentially as described in (35). The equivalent 0.35 OD_600_ units of spores, or ∼1 x 10^7^ spores, were suspended in 500 µL spore lysis buffer (0.3 mM (NH_4_)_2_SO_4_, 6.6 mM KH_2_PO_4_, 15 mM NaCl, 59.5 mM NaHCO_3_, 35.2 mM Na_2_HPO_4_). Spores in lysis buffer were split into two tubes, with one set being incubated at 37°C for 1 hour (untreated control) and the other set at 95°C for 1 hour. Samples were spun at 15,000 rpm for 2 minutes. 10 µL of supernatant was added to 115 µL of 10 mM Tris, 150 mM NaCl buffer with 800 µM terbium chloride in a black 96-well plate. Boiled WT spore supernatant was added to 10 mM Tris, 150 mM NaCl buffer without terbium chloride to determine background fluorescence. The fluorescence of terbium chloride (45) was measured in a Synergy H1 microplate reader (Biotek; excitation: 270 nm, emission: 545 nm, gain: 100) after a 15 min incubation at room temperature. The relative fluorescence units (RFU) reported represent the fluorescence after the background fluorescence (WT spores, no terbium chloride) has been subtracted. Data shown are the average fluorescence measured from three independent spore purifications.

### Spore viability following heat treatment

Spore viability after heat treatment was determined essentially as described in (35). Briefly, 1.75 OD_600_ equivalents of spores, or ∼5 x 10^7^ spores, were suspended in 500 µL sterile water. Spores were aliquoted into 7 tubes and incubated at 37°C (no heat), 50°C, 60°C, 65°C, 70°C, 75°C, or 80°C for 15 minutes. After incubation, spores were serially diluted into 1x PBS and the dilutions were plated onto BHIS with taurocholate agar media. After overnight incubation, colonies were enumerated to determine the number of germinated spores. Data shown are the averages from three independent spore purifications.

### Optical density (OD_600_)-based germination assay

The germination kinetics of spores were analyzed by measuring OD_600_ of spores over time in a microplate reader similarly to what was done in (28, 30). For all OD_600_ based kinetic assays, about 0.17 OD_600_ equivalents of spores, or ∼5 x 10^6^ spores, per condition tested were suspended in BHIS media or germination buffer (50 mM HEPES, 100 mM NaCl, pH 7.5). Spores were aliquoted into wells of a clear, flat bottomed 96-well plate (Falcon) with either calcium or glycine at indicated final concentrations to a final volume of 180 µL. As indicated, 20 µL of taurocholate (final concentration 0.2% (3.8 mM), 0.5% (9.5 mM), or 1% (19 mM) w/v) was added. For the EGTA experiments, the germination buffer included 50 mM glycine, and 2 µL of CaCl_2_ and/or EGTA stock (100X solution relative to the final concentration) was included in the 20 µL taurocholate solution. The OD_600_ was measured at 37°C every 3 minutes for 1 hour with continuous linear shaking in a Synergy H1 microplate reader (Biotek). All germinant stocks solutions were solubilized in water and filter sterilized; an initial 0.5 M EGTA stock was made in 0.1 N HCl. To measure germination kinetics in response to varying concentrations of taurocholate, glycine, or calcium, germinant solutions were serially diluted in sterile water before being mixed with spores in a 96-well plate. The change in OD_600_ was calculated as the ratio of the OD_600_ at each time point to the OD_600_ at time zero. To calculate the average change in OD_600_/minute at various concentrations of taurocholate, glycine, or calcium, the period of greatest change during the germination assay was first determined. For various concentrations of taurocholate and calcium, the greatest change was between 0 and 6 minutes and for various concentrations of glycine, the greatest change was between 0 and 15 minutes. To calculate the average change in OD_600_/min, the OD_600_ from the start of this time period was subtracted from the OD_600_ at the end of the time period and divided by the time (in minutes) of the period of greatest change. Data shown for all OD_600_ based kinetic assays are averages from experiments performed on three independent spore purifications.

### SleC cleavage analysis

Spores were diluted to about 0.35 OD_600_ equivalents per time point tested (0.7 OD units total) into germination buffer (50 mM HEPES, 100 mM NaCl, pH 7.5) with 50 mM glycine. One third of the spore volume was removed as a starting time point control. Taurocholate (TA) was added to the remaining two thirds spore volume to a final concentration of 0.2% (3.8 mM) w/v. Spores were incubated at 37°C and 100 µL samples were taken at 30- and 60-minutes post TA addition. Samples were centrifuged to pellet spores and supernatant was removed. Spores were resuspended into 50 µL EBB buffer (8 M urea, 2 M thiourea, 4% (w/v) SDS, 2% (v/v) β-mercaptoethanol) for further analysis by western blotting.

### Western blot analysis

Spore samples in EBB from SleC cleavage assays were boiled for 20 minutes with vortex mixing and 4X NuPAGE LDS Sample Buffer (Thermo Fisher) was added to stain samples with bromophenol blue. Samples were boiled for an additional 5-10 minutes with vortex mixing after the addition of sample buffer. Samples were resolved by electrophoresis on a 12% SDS-PAGE gel and transferred to a Millipore-Immobilon PVDF membrane for western blot analysis. Membranes were blocked with a 1:1 dilution of Odyssey Blocking Buffer and 1X phosphate buffered saline (PBS) with 0.1% (v/v) Tween 20 and probed with rabbit polyclonal anti-CotA antibodies (50) and monoclonal mouse anti-SleC antibodies (27). The anti-CotA antibody was used at a 1:1000 dilution and the anti-SleC antibody was used at a 1:5000 dilution. IRDye 680CV and 800CW infrared dye-conjugated secondary antibodies were used at 1:20,000 dilutions. Secondary antibody fluorescence emissions were detected on an Odyssey LiCor Clx. Results shown are representative Western blot analyses from three independent spore isolations.

### Phase-contrast microscopy analyses of spore hydration

Spores were diluted to about 0.35 OD_600_ into germination buffer (50 mM HEPES, 100 mM NaCl, pH 7.5). Spores were further diluted 1:5 in sterile water and 1.5 µL was added to an agarose pad on a microscope slide and covered with a cover slip for analysis by phase-contrast microscopy. Agarose pads were made by adhering a Gene Frame (ThermoFisher Scientific) to a glass slide and filling the Gene Frame with 1%, 19 mM (w/v) agarose in germination buffer. Agarose pads were incubated at 4°C to solidify. Spores were analyzed on an Axioskop (Carl Zeiss, West-Germany) upright microscope, a 100x 1.3 NA Plan NeoFluar phase contrast objective, and images were taken using a Hamamatsu C4742-95 Orca 100 CCD Camera with a 25 ms exposure. Images were taken of fields with well-separated spores to ensure better identification of spores in downstream analysis. Phase-contrast images were analyzed using the MicrobeJ plug-in (44) for ImageJ to identify individual spores and measure the intensity of phase-bright spore core. MicrobeJ analysis was completed with the following parameters: width [p]: 0.6-1, circularity [0-1]: 0.5-1, curvature [0-max]: 0-0.4, sinuosity [0-max]:0-1.1, angularity [rad]: 0-0.5, solidity [0-max]: 0-max, intensity [0-max]: 0-200, Z-score: 0-max, feature [0-max]: 0-max, poles [0-max]: 0-max, range [p]: 0-max, symmetry: 0-max, holes [0-max]: 0-0. The percent of spores correctly identified in an image varied between images and *C. difficile* strains, but at least 275 spores were analyzed per strain in a replicate experiment. The average intensity (AU) of individual spores was graphed as a proxy for spore hydration. The data shown derive from three independent spore isolations.

### Single spore germination analyses

#### PySpore Validation Experiments

To generate agarose pads for the analyses, 2X BHIS containing 1%, 19 mM TA was mixed with molten 3.5% (w/v) Top-Vision Low Melting Point Agarose in sterile H_2_O to a final concentration of 1.75% agarose and 0.5% (9.5 mM) TA. The resulting media was added to a 125 μL gene-frame (Thermo-Fisher Scientific) adhered to a clean glass microscope slide and solidified. WT, Δ*pdaA,* and Δ*cwlD* spores were diluted to 0.2-0.5 OD_600_ in sterile H_2_O in biological triplicate and 1μl was spotted on to a #1.5 coverslip and allowed to dry at 37°C. After the spores dried, the coverslip was inverted onto the agar pad and imaged immediately imaged via time-lapse phase-contrast microscopy at 37°C.

#### Glycine and Calcium Experiments

Agarose pads were made containing germination buffer (50 mM HEPES, 100 mM NaCl, pH 7.5), 1.75% (w/v) Top-Vision Low Melting Point Agarose, 0.2% (3.8 mM) TA (w/v), and either 10 mM Gly or 30 mM CaCl_2._ The resulting media was added to a 125μl gene-frame (Thermo-Fisher Scientific) adhered to a clean glass microscope slide and solidified. WT, Δ*dpaAB* spores were diluted to 0.2-0.5 OD_600_ in sterile H_2_O in biological triplicate and 1μl was spotted on to a #1.5 coverslip and allowed to dry at 37°C. After the spores dried, the coverslip was inverted onto the agar pad and imaged immediately imaged via time-lapse phase-contrast microscopy at 37°C.

#### Spore Exudate Experiments

To generate the germination exudates used to make the agarose pads for analyzing spore germination using time-lapse microscopy, the equivalent of 0.7 OD_600_ units were diluted into germination buffer (50 mM HEPES, 100 mM NaCl, pH 7.5). The TA germinant and glycine co-germinant were added to final concentrations of 0.5%, 9.5 mM TA and 2.5 mM glycine and incubated at 37°C for 4 hours. The germination exudate was harvested from the spores by centrifuging the samples at 15,000 rpm for 2 minutes to pellet the spores and removing the supernatant (i.e. the germination exudate). The supernatant/ exudate was centrifuged again at 15,000 rpm for 2 minutes. If the germination exudate was not immediately used, it was stored at 4°C. Otherwise, the supernatant was incubated for ∼10 minutes at 37°C and then 200 µL of the exudate was added to 100 µL of melted 3.3% agarose in germination buffer (50 mM HEPES, 100 mM NaCl, pH 7.5) for final concentration of 1% agarose. For the Ca^2+^ “add-back” experiment, CaCl_2_ added to a final concentration of 190 µM during the pre-warming step at 37°C. The agarose-germination exudate solution was gently vortexed and then added to a Gene Frame adhered to a microscope slide. The agarose pad was incubated at 4°C to solidify the agarose pad during which time 0.175 OD_600_ equivalents of WT spores were diluted 1:4 in sterile water and vigorously pipetted to break up spore clumps. 1.5 µL of this WT spore mixture were added to agarose pad; the pad was then sealed with a cover slip and immediately analyzed by phase-contrast time-lapse microscopy.

#### Data Analysis

Resulting image stacks were registered if significant drift of the FOV was encountered, segmented, and quantified using a custom GUI based Python application developed in-house called PySpore. PySpore utilizes phase cross correlation for image registration/alignment of the image stack. To segment spores a black-hat transform is applied to each image to enhance the dark features of the spore, namely the spore coat, while also eliminating background and correcting for discrepancies in illumination over the field. A threshold is chosen from the black-hat processed image and a segmentation mask is generated. Each ROI then undergoes a simple pixel overlap based image processing algorithm to track its position throughout the movie and correct any discrepancies in spore ID, and measurements are extracted including size, shape, and intensity from the phase-contrast images and are output as a .csv file for downstream analysis.

All downstream analyses including statistical tests were done in Python. Spore clumps were excluded based on area; spores tracked for less than 50 frames were also excluded from further analysis. Additionally, highly noisy traces were excluded after evaluating the coefficient of variation (CV) at multiple windows for each time series and discarded based on a threshold CV value and the number of times that was exceeded in the time series.

Germination rate and time to germination were quantified from spores where germination was tracked (>20% decrease in intensity from start to end). Maximal germination rate was determined by evaluating the first derivative of 1-D smoothing spline fit to the data at each time point. Time to germination was defined as the time point when the max germination rate is reached.

PySpore code is available here: https://github.com/jribis89/PySpore

#### Microscope Hardware

Images were taken every 30 seconds using a Leica DMi8 inverted microscope equipped with a 63x 1.4NA Plan Apochromat phase-contrast objective, Leica Adaptive Focus Control (AFC), a high precision motorized stage (Pecon), an incubator (Pecon), and a Leica DFC9000-GTC sCMOS camera for all time-lapse analyses.

## Supplementary Figure Legends

**Figure S1. CaDPA mutant spores are less “phase-bright”.**

**Figure S2. Spore purification efficiency and heat resistance of CaDPA mutant spores. Figure S3. Spline determination of germination rate and germination time.**

**Figure S4. Comparison of PySpore and SporeTrackerX.**

**Figure S5. Comparison of CaDPA mutant spore germination in the presence of glycine vs. Ca^2+^ co-germinant.**

**Figure S6. Micromolar concentrations of Ca^2+^ appear to be required for *C. difficile* germination to proceed**

