## Supplementary files for "Single-spore germination analyses reveal that calcium released during *Clostridioides difficile* germination functions in a feed-forward loop"

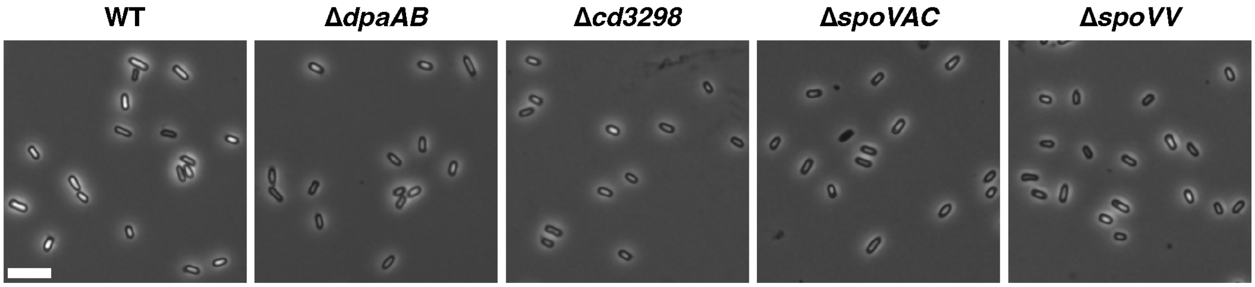


**Figure S1. CaDPA mutant spores are less “phase-bright”.** Example phase-contrast images used to determine the intensities shown in Fig 1C. Scale bar = **5 μm.**

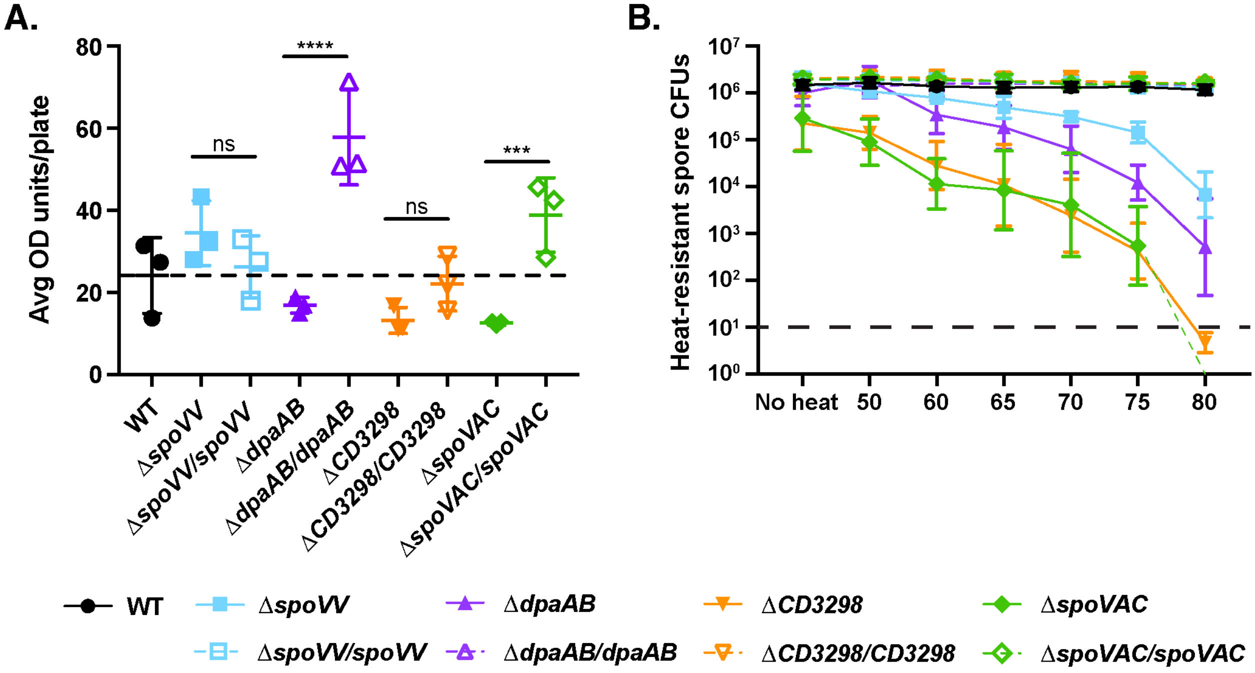


**Figure S2. Spore purification efficiency and heat resistance of CaDPA mutant spores.** (A) Average spore yield obtained from purifications of the indicated strains based on three biological replicates. Yields were determined by measuring the optical density of spore purifications at 600 nm and are expressed as $OD_{600}$ units obtained per plate. Statistical significance was determined using one-way ANOVA and Tukey's test.(B) Heat resistance of purified spores from the indicated strains. Spores were incubated for 15 min at the indicated temperature, plated on rich media containing TA germinant, and the number of CFUs produced by germinated spores was enumerated.

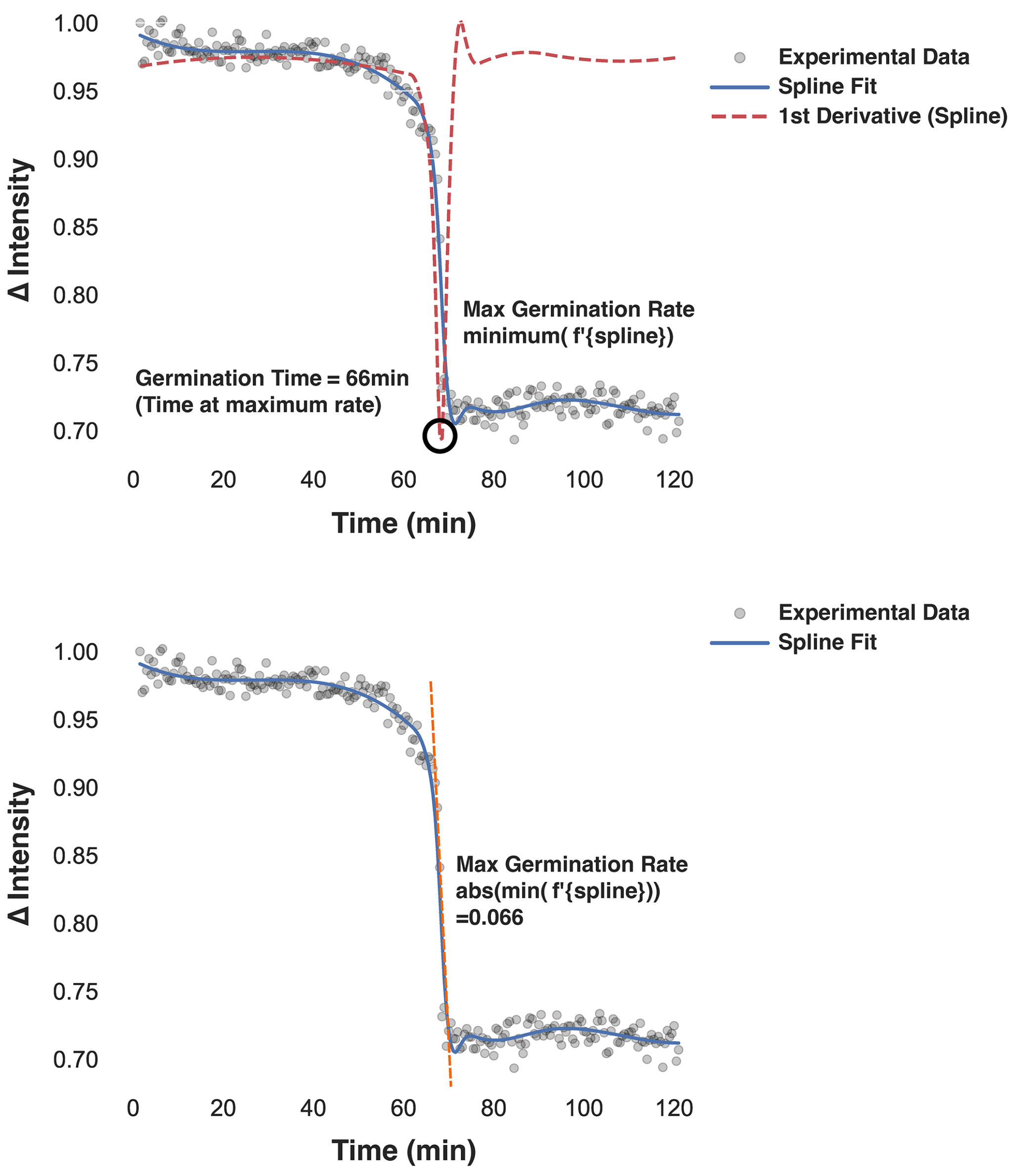


**Figure S3. Spline determination of germination rate and germination time.** Plots showing change in intensity over time. Points indicate raw data, solid blue line shows fit of 1D smoothing spline. The dashed red line shows the first derivative of the fitted spline evaluated at each timepoint, and the minimum is circled to highlight the point where maximal germination rate is reached, which is the time to germination. The bottom panel includes the line tangent to the point of maximal germination rate, shown in orange. Max germination rate is conveyed as an absolute value for clarity.

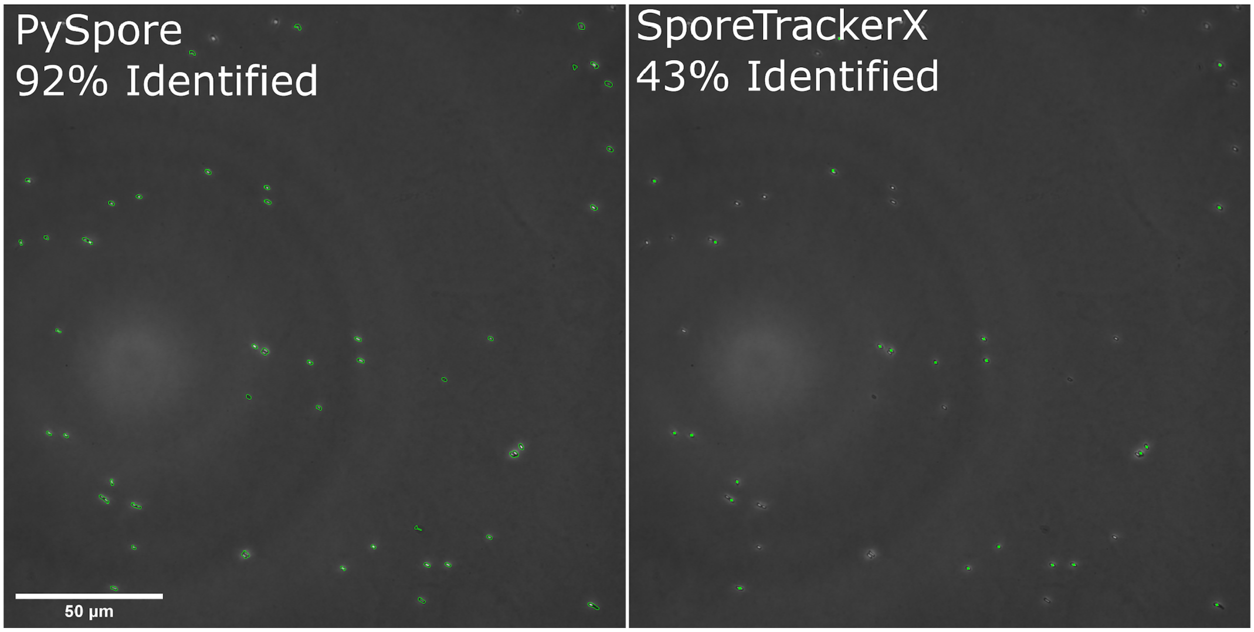


**Figure S4. Comparison of PySpore and SporeTrackerX.** Single frame from movie of WT *C.difficile* germinating on a 1.5% agarose pad containing BHIS and 0.5% sodium taurocholate. Spores in the image were segmented with both PySpore and SporeTrackerX and the percentage of spores successfully identified in the field was calculated.

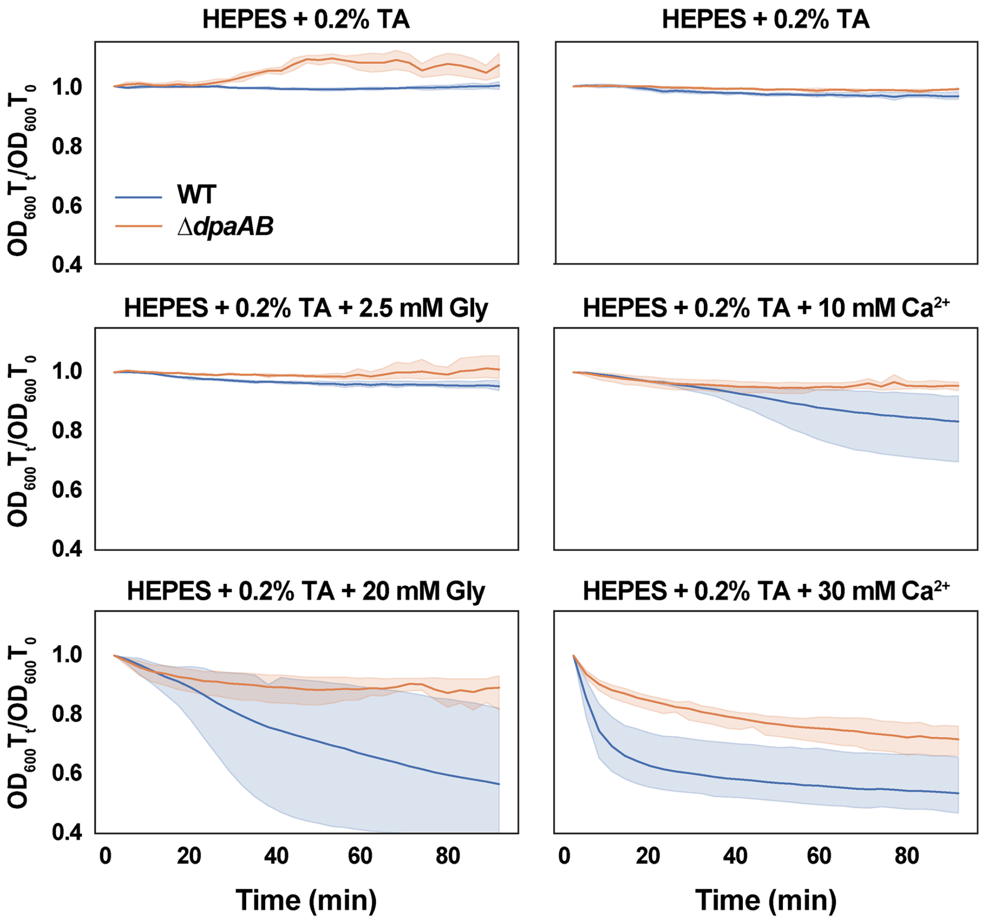


**Figure S5. Comparison of CaDPA mutant spore germination in the presence of glycine vs. $Ca^{2+}$ co-germinant**. Optical density ($OD_{600}$)-based analyses of spore germination over time in the presence of the indicated concentrations of taurocholate (TA) and either glycine (Gly) or $Ca^{2+}$. The ratio of the $OD_{600}$ of each strain at a given time point relative to at time zero was plotted. Error bands are the standard error.

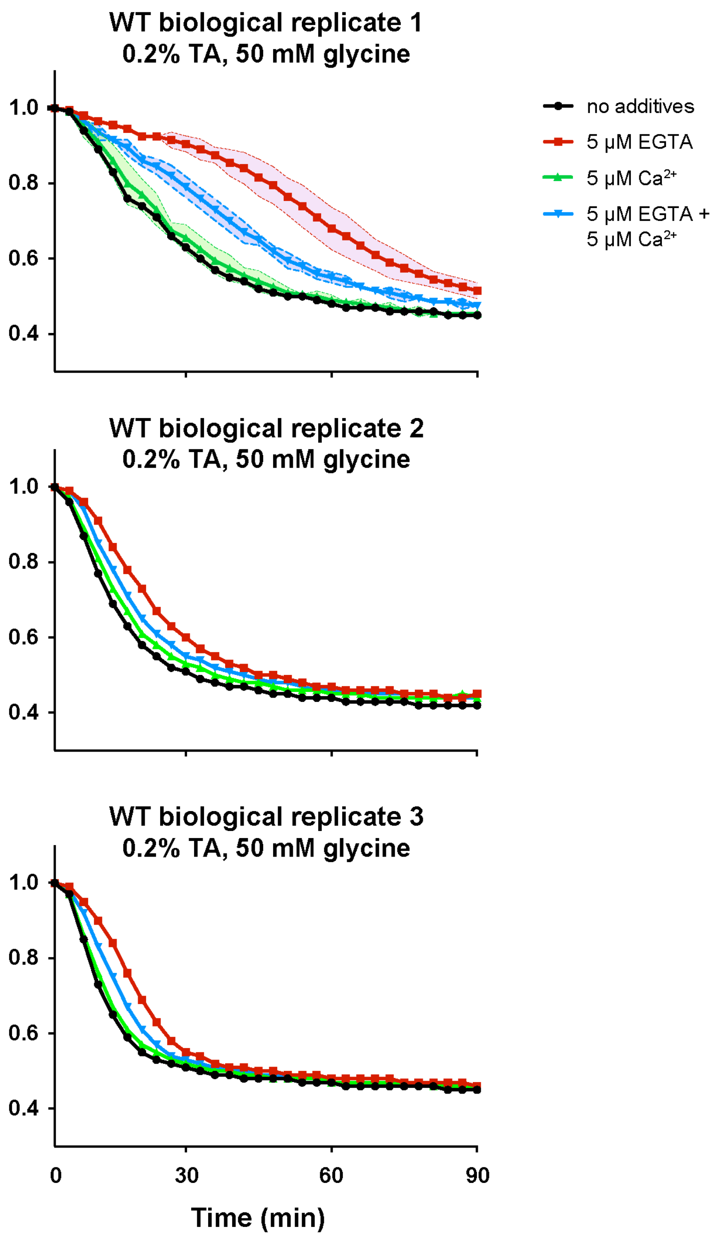


**Figure S6. Micromolar concentrations of $Ca^{2+}$ appear to be required for *C. difficile* germination to proceed.** Optical density ($OD_{600}$)-based analyses of WT spore germination over time in the presence of 0.2% (3.8 mM) taurocholate and 50 mM glycine and the indicated concentrations of EGTA (A) and 5 µM EGTA and/or 5 µM $Ca^{2+}$. The ratio of the $OD_{600}$ at a given time point relative to at time zero was plotted. Three independent experiments (conducted on three independent spore purifications) are shown. The shaded error bars indicate the standard deviation for each time point measured for three technical replicates of the first biological replicate of spores analyzed. Due to variability in the behavior of independent WT spore preparations, the results were not averaged, so no statistical analyses were not performed. The variability appears to derive from CaDPA release during germination, since at 5 µM $Ca^{2+}$ supplementation, this level of variability was not observed in CaDPA-less spores ($\Delta$*dpaAB*, Fig 7).
